# Optimal markers for the identification of *Colletotrichum* species

**DOI:** 10.1101/659177

**Authors:** Willie Anderson dos Santos Vieira, Priscila Alves Bezerra, Anthony Carlos da Silva, Josiene Silva Veloso, Marcos Paz Saraiva Câmara, Vinson Patrick Doyle

**Affiliations:** Universidade Federal Rural de Pernambuco – UFRPE. Departamento de Agronomia. Rua Manuel de Medeiros, s/n - Dois Irmãos, Recife – Pernambuco, 52171-900; Department of Plant Pathology and Crop Physiology, Louisiana State University – LSU, AgCenter, Baton Rouge, Louisiana, United States of America, 70808

**Keywords:** Accuracy, Anthracnose, Barcoding, Phylogenetic informativeness, Standardization

## Abstract

*Colletotrichum* is among the most important genera of fungal plant pathogens. Molecular phylogenetic studies over the last decade have resulted in a much better understanding of the evolutionary relationships and species boundaries within the genus. There are now approximately 200 species accepted, most of which are distributed among 13 species complexes. Given their prominence on agricultural crops around the world, rapid identification of a large collection of *Colletotrichum* isolates is routinely needed by plant pathologists, regulatory officials, and fungal biologists. However, there is no agreement on the best molecular markers to discriminate species in each species complex. Here we calculate the barcode gap distance and intra/inter-specific distance overlap to evaluate each of the most commonly applied molecular markers for their utility as a barcode for species identification. Glyceraldehyde-3-phosphate dehydrogenase (GAPDH), histone-3 (HIS3), DNA lyase (APN2), intergenic spacer between DNA lyase and the mating-type locus *MAT*1-2-1 (APN2/MAT-IGS), and intergenic spacer between GAPDH and a hypothetical protein (GAP2-IGS) have the properties of good barcodes, whereas sequences of actin (ACT), chitin synthase (CHS-1) and nuclear rDNA internal transcribed spacers (nrITS) are not able to distinguish most species. Finally, we assessed the utility of these markers for phylogenetic studies using phylogenetic informativeness profiling, the genealogical sorting index (GSI), and Bayesian concordance analyses (BCA). Although GAPDH, HIS3 and β-tubulin (TUB2) were frequently among the best markers, there was not a single set of markers that were best for all species complexes. Eliminating markers with low phylogenetic signal tends to decrease uncertainty in the topology, regardless of species complex, and leads to a larger proportion of markers that support each lineage in the Bayesian concordance analyses. Finally, we reconstruct the phylogeny of each species complex using a minimal set of phylogenetic markers with the strongest phylogenetic signal and find the majority of species are strongly supported as monophyletic.

## 1. INTRODUCTION

*Colletotrichum* is among the largest groups of phytopathogenic fungi and includes the causal agents of anthracnose and other diseases on seeds, stems, leaves and fruits of important temperate and tropical crops (Cai et al., 2009, Cannon et al., 2012). It is also among the most common genera of endophytic fungi, fungi that live within plant organs without producing any symptoms of disease (Cannon et al., 2012). Due to its economic and scientific importance, *Colletotrichum* was ranked as the eighth most important phytopathogenic fungus in the world by plant pathologists (Dean et al., 2012).

Species identification is necessary to understand disease epidemiology and develop strategies to control the disease successfully (Cai et al., 2009). However, *Colletotrichum* taxonomy and systematics has been a challenge since the genus was introduced by Corda (1831). *Colletotrichum* species were historically circumscribed on the basis of phenotypic features and a strong emphasis on the host species from which the specimens were isolated under the assumption of host specificity, which led to more than 900 species being recognized until revisionary work more than a century after its introduction (von Arx, 1957; Sutton, 1980). *Colletotrichum* identification based on morphological characters is problematic due to plasticity and variation induced by experimental conditions (Vieira et al. 2017), and all life stages are not frequently produced in culture (Samarakoon et al., 2018). The absence of stable phenotypic characters has limited our understanding of phylogenetic relationships within *Colletotrichum* and made the recognition of species boundaries unreliable and confusing (Cai et al., 2009). To address this problem, Cai et al. (2009) proposed a guideline for *Colletotrichum* species recognition based on a polyphasic approach, which comprises the use of cultural, morphological, physiological and pathogenicity characters in combination with phylogenetic analysis of nucleic acid sequences.

The earliest phylogenetic studies of *Colletotrichum* using DNA sequences were published by Mills et al. (1992) and Sreenivasaprasad et al. (1992). Polymorphisms in the ITS1 region of the nrDNA were used to distinguish *Colletotrichum* species. However, while the nrITS region is the most widely sequenced region and has been chosen as the barcode locus for the Fungi, the utility of this region is limited for systematic studies in *Colletotrichum*. Species diversity is usually underestimated when based on nrITS sequences alone (Crouch et al., 2009a) and it has been demonstrated to have little phylogenetic utility (Doyle et al. 2013; Vieira et al. 2017). However, several additional markers have been applied for multilocus phylogenetic inference to resolve the boundaries of cryptic species in the genus (Cai et al. 2009; Damm et al. 2009; Doyle et al., 2013; Hyde et al., 2009; Lima et al., 2013; Liu et al., 2016; Samarakoon et al., 2018; Veloso et al., 2018; Vieira et al., 2014, 2017).

According to the most recent synopsis of the genus published in 2017, 188 *Colletotrichum* species have been described and incorporated into molecular phylogenies. Among these species, 164 were distributed among 11 species complexes and an additional 24 species had not been assigned to a species complex (Marin-Felix et al., 2017). Additional species were recently described and three additional clades were declared to represent new species complexes (Cao et al., 2018; Damm et al., 2019; Samarakoon et al., 2018). Due to its global distribution and ecological and economic importance, research groups around the world are working concomitantly to address regional diversity. However, the set of phylogenetic markers used to discriminate species is variable by species complex and no standard set of markers has been adopted based on objective criteria (Marin-Felix et al., 2017), making it difficult to combine data from disparate research groups and reliably infer phylogenies and delimit species boundaries. Currently, thirteen different molecular markers are commonly sequenced among the various *Colletotrichum* species complexes: actin (ACT), DNA lyase (APN2), intergenic spacer between DNA lyase and the mating-type locus *MAT*1-2-1 (APN2/MAT-IGS), calmodulin (CAL), chitin synthase (CHS-1), glyceraldehyde-3-phosphate dehydrogenase (GAPDH), intergenic spacer between GAPDH and a hypothetical protein (GAP2-IGS), glutamine synthetase (GS), histone 3 (HIS3), nuclear rDNA internal transcribed spacers (nrITS), mating type gene (MAT1-2-1), manganese-superoxide dismutase (SOD2), and β-tubulin (TUB2).

It is known that the efficiency of PCR amplification and the distribution of phylogenetic informativeness of a given marker varies among species complexes (Hyde et al., 2013). Most studies on the utility and reliability of individual markers come from the *Colletotrichum gloeosporioides* complex (Cai et al., 2009; Liu et al., 2015; Sharma et al. 2013, Silva et al., 2012;2015; Vieira et al. 2017), and while the *C. gloeosporioides* complex has been exhaustively studied in recent years, recommendations on marker choice seem to be largely ignored. A classic case is the utility of APN2/MAT-IGS, the most powerful marker to discriminate species within the *C. gloeosporioides* complex: several recent studies described novel species within *C. gloeosporioides* complex while excluding data from APN2/MAT-IGS (Costa et al., 2018; Diao et al. 2017; Fu et al., 2019; Jayawardena et al., 2016; Oliveira et al., 2018; Sharma et al., 2017; Silva et al., 2017; Sousa et al., 2018; Wang et al., 2019). As mentioned above, this limits our ability to combine data from regional studies to develop an accurate understanding of global diversity and phylogenetic relationships within the genus.

While robust phylogenetic inference and reliable species delimitation relies on the use of quality markers, studies on the performance of different molecular markers for phylogenetic inference are missing for the majority of *Colletotrichum* species complexes. In addition to the challenges discussed above, this also presents practical problems for plant pathologists and ecologists who are looking to reliably identify a large collection of isolates using molecular data. It is impractical for researchers to sequence several loci, many of which may be of little phylogenetic utility, across several hundred isolates simply for species identification. The aim of the present study was to evaluate the phylogenetic informativeness of different molecular markers used in *Colletotrichum* systematics and determine the optimal set of markers for each species complex. From this, we hope to establish a consensus on the minimal set of markers that can be used for *Colletotrichum* species identification and delimitation and provide a practical reference for the large community of researchers working on developing a better understanding of global diversity, life history, and ecology of the genus.

## 2. MATERIAL AND METHODS

### 2.1 Datasets

We compiled several datasets to analyze the barcoding utility, phylogenetic signal, and genealogical concordance of ACT, APN2, APN2/MAT-IGS, CAL, CHS-1, GAPDH, GAP2-IGS, GS, HIS3, nrITS, and TUB2 for all *Colletotrichum* species complexes described to date except for the *C. caudatum* complex, which was excluded because nrITS was the only marker available for all species. These datasets were compiled from published sequences retrieved from GenBank (Supplementary File S1). Since testing the accuracy of prior species delimitations were not the main focus of this study, we assumed that species boundaries established in previous studies were accurate.

Thirteen species complexes were investigated in our study: *Colletotrichum acutatum, C. boninense, C. dematium, C. destructivum, C. dracaenophilum, C. gigasporum, C. gloeosporioides, C. graminicola, C. magnum, C. orbiculare, C. orchidearum, C. spaethianum* and *C. truncatum*. Some species within each complex were not included in the alignments due to the absence of sequences for several markers since some of the analyses employed in the present work do not allow missing data. Some markers were not analyzed due to a small number of species or isolates with sequences available. The inclusion of these markers will drastically reduce the number of species that can be included in each set of analyses (e.g. GAPDH, HIS3, APN2 and APN2/MAT-IGS in the *C. graminicola* species complex).

### 2.2 Multiple sequence alignment

Multiple sequence alignments (MSA) of each locus were estimated individually for each species complex. Sequences were compiled using the GenBank tool implemented in MEGA 7 (Kumar et al., 2016). MSAs were estimated with the online version of MAFFT 7 (Katoh et al., 2002; Katoh & Standley, 2013) using the G-INS-i iterative refinement method and the 200PAM / k=2 nucleotide scoring matrix. External gaps were trimmed in MEGA 7 before uploading MSAs to the GUIDANCE2 server (http://guidance.tau.ac.il/ver2/) (Sela et al., 2015) to access the alignment confidence scores under the following parameters: MAFFT as the MSA algorithm; max-iterate=0; pairwise alignment method=6mer; 100 bootstrap replicates. Unreliable alignment regions were filtered by masking residues with scores below the lowest cutoff, as proposed by Vieira et al. (2017). MSAs were converted to nexus format, concatenated, and partitioned into a multilocus matrix using SequenceMatrix 1.8 (Vaidya et al., 2011). The number of invariable (I), variable (V), singletons (S) and parsimony informative (PI) characters of the single locus alignments were calculated using DnaSP 5.10 (Librado and Rozas,2009).

### 2.3 DNA barcoding

The effectiveness of markers to discriminate species within each species complex was assessed by the barcode gap distance and intra/inter-specific distance overlap (Hebert et al. 2003). Intra- and inter-specific distances were calculated for each single locus alignment in MEGA 7. Single isolate species were removed from alignments. Distances were calculated under the Kimura-2-parameter model, allowing for substitution rates to differ among transitions and transversions, uniform rates among sites, and gaps treated as pairwise deletions. Distance values were sorted in Microsoft Excel Professional Plus 2016 and summary statistics were calculated (maximum, minimum and mean distance). The barcode gap was represented by the difference between the mean interspecific and intraspecific distances (Hebert et al. 2003). The distance overlap percentage represents how much the intraspecific distance overlaps with the interspecific distance and was calculated as follows: max intraspecific distance ÷ max interspecific distance × 100. Markers useful as barcodes will have a large barcode gap and a small intra/inter-specific distance overlap.

### 2.4 Assessment of phylogenetic informativeness

The phylogenetic informativeness of markers commonly employed in *Colletotrichum* systematics was estimated using the application PHYDESIGN (Lopez-Giraldez and Townsend, 2011). Maximum likelihood (ML) trees were inferred for each species complex using the concatenated alignments reduced to a single representative isolate per species. Phylogenies were estimated in RAxML - HPC2 (Stawatakis, 2014) implemented on CIPRES Science Gateway portal (https://www.phylo.org/portal2/home.action). ML tree searches were done assuming the GTRGAMMA model and bootstrap support calculated with 1000 pseudoreplicates (-m GTRGAMMA -p 12345 -k -f a -N 1000 -x 12345). ML trees were converted to rooted ultrametric trees using the ‘chronos’ function in the ape package (Paradis et al., 2004) using R Studio 1.1.442 (R Core Team, 2017). Trees were calibrated with an arbitrary time scale with time=0 at the tips and time=1 at the root. The ultrametric trees and the corresponding partitioned alignment were used as input files in PHYDESIGN and the substitution rates were calculated using the program HyPhy (Pond et al., 2005). The substitution rates estimated by the maximum likelihood algorithm used by HyPhy are nonsensical for some sites in the alignment resulting in very recent ‘phantom’ peaks that have no biological meaning. These peaks are likely the result of indels or ambiguous sites in the alignment, therefore these alignment positions with poorly estimated substitution rates were excluded from some genes prior to phylogenetic informativeness profiling at the recommendation of the authors of PhyDesign (http://phydesign.townsend.yale.edu/faq.html). Since the markers included in this study and in most systematic studies of *Colletotrichum* do not require more than two sequencing reads to sequence the full length of the locus (representing the same sequencing effort and cost), the phylogenetic informativeness values (PIV) were calculated on a net basis. The variable PImax represents the time in which a given marker reaches the maximum PIV and was used to determine the divergence time in which the marker is most informative (Fong and Fujita, 2011). We ranked the markers according to the PIV values and the usefulness of markers was assessed through the profile shape: low and flat curves represent the least informative markers; high and sharp peaks represent the most informative markers.

The percentage of markers that resolve a given species was estimated using a Bayesian Concordance Analysis – BCA (Ané et al., 2007; Larget et al., 2010). Although BCA is a coalescent-based method to estimate species trees (Ané et al., 2007; Baum, 2007; Larget et al., 2010), this methodology can also be used to quantify the proportion of markers that support a given clade, which is represented by the concordance factor (CF). Individual locus trees were inferred in MrBayes 3.2.6 (Ronquist et al., 2012) implemented on the CIPRES cluster with four runs, each run with four Markov chain Monte Carlo (MCMC) chains run for 10,000,000 generations, sampled every 5,000 generations, totaling 2,001 trees per run. The frequency of distinct topologies in the posterior distribution were summarized using mbsum distributed with BUCKy 1.4.4 (Ané et al., 2007; Larget et al., 2010) skipping the first 25% of the trees as burn-in (-n 501). Primary concordance trees were estimated using bucky and tree summary files output by mbsum were used as input files. Concordance analyses were performed with a discordance factor (α) set at 1, four MCMC chains, 1,000,000 generations, and the first 25% generations were discarded as burn-in (-a 1 -k 4 -n 1000000 -c 4 -s1 23546 -s2 4564).

The Genealogical Sorting Index (GSI) was employed to identify the markers that recover each species as monophyletic. GSI is an objective method that infers the proportion of input trees for which a clade (species as applied here) are found to be monophyletic and if the observed monophyly is greater than would be observed by chance given the size of the data matrix (GSI=1 indicates monophyly) (Cummings et al., 2008; Sakalidis et al., 2011), and can also be used to compare individual markers according to their ability to discriminate species (Doyle et al., 2013). This methodology can be applied to phylogenies inferred from a single locus as well as multilocus analysis (Sakalidis et al., 2011). ML analyses were performed using the single and multi-locus concatenated alignments. Species with a single isolate were removed from the alignments. Analyses were carried out in RAxML as described above with the number of pseudoreplicates reduced to 100 and outgroup isolates specified prior to analysis. Rooted bootstrap trees were used as input files and each tip was assigned to a species. GSIs were calculated using the GSI.py script and *P*-value estimated from 100 permutations of each dataset (Cummings et al., 2008). The GSI values were converted to heatmaps in the Heatmapper web server (http://www.heatmapper.ca) to aid in visualization (Babicki et al., 2016).

### 2.5 Selection of best minimal sets of markers

The selection of a minimal set of optimal markers for robust phylogenetic inference within each *Colletotrichum* species complex was based on results from the phylogenetic informativeness profiling and GSIs. The markers were selected according to the following criteria:

1. A minimum of three markers per complex based on ranking according to PIV were selected. Three independent markers allow for the application of the genealogical concordance phylogenetic species recognition criteria – GCPSR (Dettman et al., 2003, Taylor et al. 2000), a commonly applied set of criteria for phylogenetic species recognition in fungal systematics.
2. All species must be recognized as monophyletic by at least one of the selected markers. GSIs were checked to confirm if each species in the complex is recovered as monophyletic (GSI = 1) by at least one of the selected markers.
3. ML trees were inferred from concatenated alignments of the three best markers; if some species clade was poorly supported and/or all species were not recovered as in the multilocus analyses with all markers (unresolved relationships/polytomy), other markers were progressively concatenated in decreasing order of phylogenetic informativeness until all species were well resolved.

Once the best markers were chosen, the GSI was calculated for this phylogeny to determine the level of species monophyly when only the best markers are concatenated. The BCAs were also performed with this dataset to elucidate if the species could be recognized by the majority of the selected markers.

## 3. RESULTS AND DISCUSSION

### 3.1 Alignment statistics

GAPDH, HIS3, and TUB2 were the most variable markers in the majority of *Colletotrichum* species complexes, with PI characters ranging from 10—109, 11—82, and 12—114, respectively (Table 1). PI characters for GS ranged from 63 to 93 and it was the most variable marker within the *C. gigasporum* and *C. orbiculare* species complexes. The APN2/MAT-IGS and GAP2-IGS, which are employed only in the *C. gloeosporioides* species complex, had 192 and 115 PI characters, respectively, and were the most variable markers within this complex. In contrast, nrITS presented the fewest PI characters for most species complexes (0—36), followed by CHS-1 (3—45) and ACT (3—63).

**Table 1.**
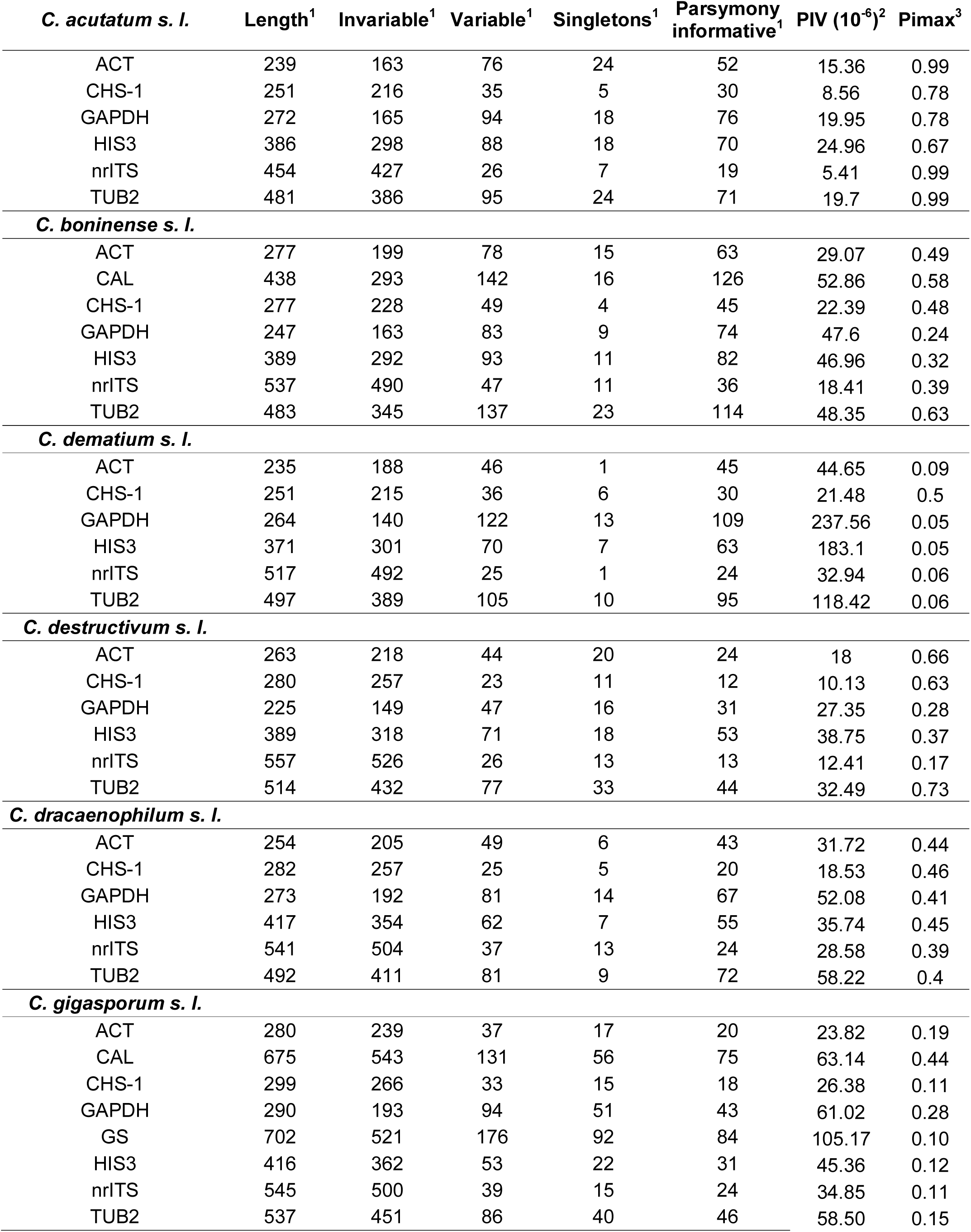
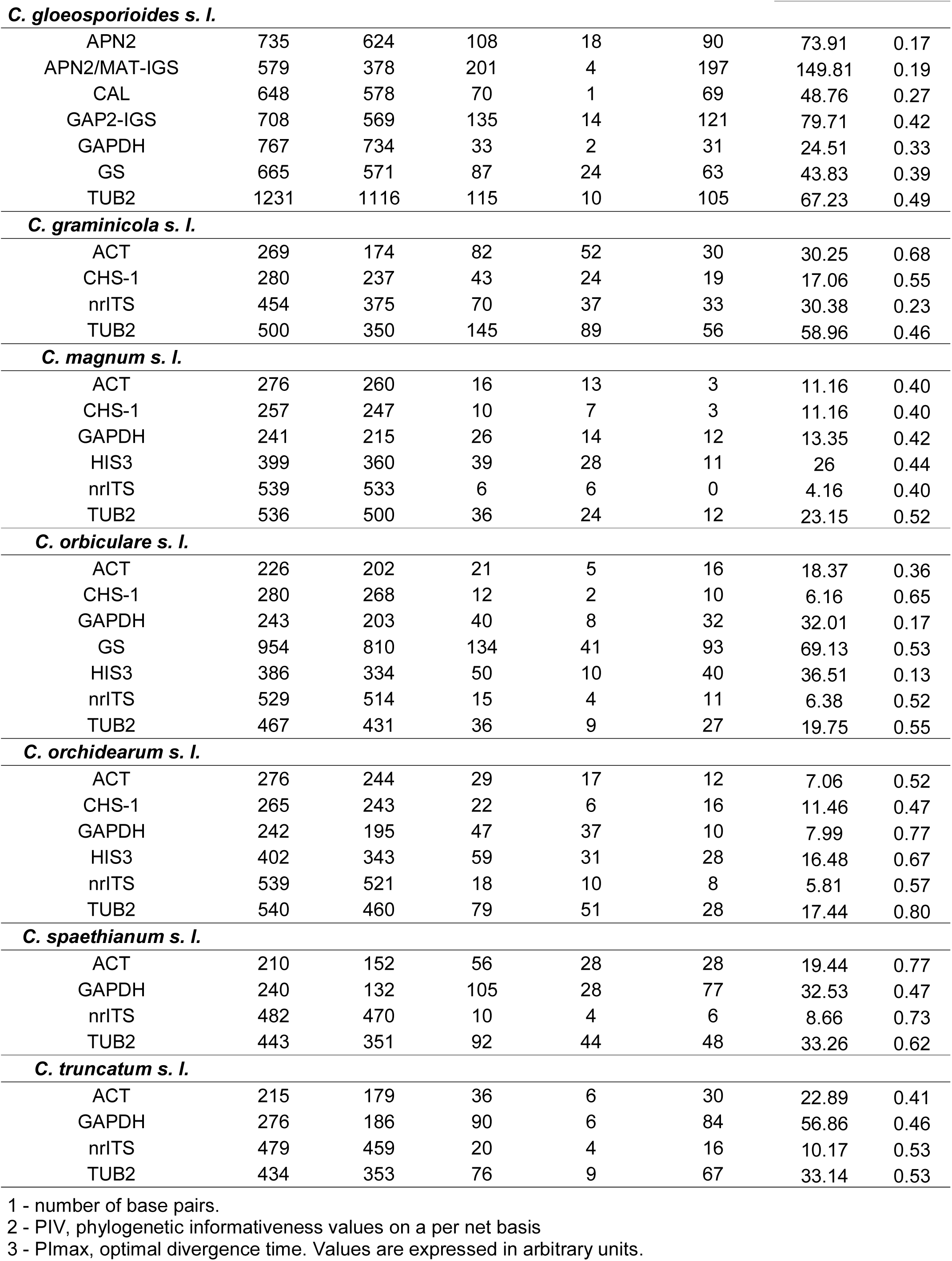
Alignment and phylogenetic informativeness profile statistics for markers used in Colletotrichum by species complex.

| *C. acutatum* s. l. | Length<sup>1</sup> | Invariable<sup>1</sup> | Variable<sup>1</sup> | Singletons<sup>1</sup> | Parsimony informative<sup>1</sup> | PIV (10<sup>-6</sup>)<sup>2</sup> | Pimax<sup>3</sup> |
| --- | --- | --- | --- | --- | --- | --- | --- |
| ACT | 239 | 163 | 76 | 24 | 52 | 15.36 | 0.99 |
| CHS-1 | 251 | 216 | 35 | 5 | 30 | 8.56 | 0.78 |
| GAPDH | 272 | 165 | 94 | 18 | 76 | 19.95 | 0.78 |
| HIS3 | 386 | 298 | 88 | 18 | 70 | 24.96 | 0.67 |
| nrITS | 454 | 427 | 26 | 7 | 19 | 5.41 | 0.99 |
| TUB2 | 481 | 386 | 95 | 24 | 71 | 19.7 | 0.99 |
| *C. boninense* s. l. |  |  |  |  |  |  |  |
| ACT | 277 | 199 | 78 | 15 | 63 | 29.07 | 0.49 |
| CAL | 438 | 293 | 142 | 16 | 126 | 52.86 | 0.58 |
| CHS-1 | 277 | 228 | 49 | 4 | 45 | 22.39 | 0.48 |
| GAPDH | 247 | 163 | 83 | 9 | 74 | 47.6 | 0.24 |
| HIS3 | 389 | 292 | 93 | 11 | 82 | 46.96 | 0.32 |
| nrITS | 537 | 490 | 47 | 11 | 36 | 18.41 | 0.39 |
| TUB2 | 483 | 345 | 137 | 23 | 114 | 48.35 | 0.63 |
| *C. dematium* s. l. |  |  |  |  |  |  |  |
| ACT | 235 | 188 | 46 | 1 | 45 | 44.65 | 0.09 |
| CHS-1 | 251 | 215 | 36 | 6 | 30 | 21.48 | 0.5 |
| GAPDH | 264 | 140 | 122 | 13 | 109 | 237.56 | 0.05 |
| HIS3 | 371 | 301 | 70 | 7 | 63 | 183.1 | 0.05 |
| nrITS | 517 | 492 | 25 | 1 | 24 | 32.94 | 0.06 |
| TUB2 | 497 | 389 | 105 | 10 | 95 | 118.42 | 0.06 |
| *C. destructivum* s. l. |  |  |  |  |  |  |  |
| ACT | 263 | 218 | 44 | 20 | 24 | 18 | 0.66 |
| CHS-1 | 280 | 257 | 23 | 11 | 12 | 10.13 | 0.63 |
| GAPDH | 225 | 149 | 47 | 16 | 31 | 27.35 | 0.28 |
| HIS3 | 389 | 318 | 71 | 18 | 53 | 38.75 | 0.37 |
| nrITS | 557 | 526 | 26 | 13 | 13 | 12.41 | 0.17 |
| TUB2 | 514 | 432 | 77 | 33 | 44 | 32.49 | 0.73 |
| *C. dracaenophilum* s. l. |  |  |  |  |  |  |  |
| ACT | 254 | 205 | 49 | 6 | 43 | 31.72 | 0.44 |
| CHS-1 | 282 | 257 | 25 | 5 | 20 | 18.53 | 0.46 |
| GAPDH | 273 | 192 | 81 | 14 | 67 | 52.08 | 0.41 |
| HIS3 | 417 | 354 | 62 | 7 | 55 | 35.74 | 0.45 |
| nrITS | 541 | 504 | 37 | 13 | 24 | 28.58 | 0.39 |
| TUB2 | 492 | 411 | 81 | 9 | 72 | 58.22 | 0.4 |
| *C. gigasporum* s. l. |  |  |  |  |  |  |  |
| ACT | 280 | 239 | 37 | 17 | 20 | 23.82 | 0.19 |
| CAL | 675 | 543 | 131 | 56 | 75 | 63.14 | 0.44 |
| CHS-1 | 299 | 266 | 33 | 15 | 18 | 26.38 | 0.11 |
| GAPDH | 290 | 193 | 94 | 51 | 43 | 61.02 | 0.28 |
| GS | 702 | 521 | 176 | 92 | 84 | 105.17 | 0.10 |
| HIS3 | 416 | 362 | 53 | 22 | 31 | 45.36 | 0.12 |
| nrITS | 545 | 500 | 39 | 15 | 24 | 34.85 | 0.11 |
| TUB2 | 537 | 451 | 86 | 40 | 46 | 58.50 | 0.15 |
| ***C. gloeosporioides s. l.*** |  |  |  |  |  |  |  |
| APN2 | 735 | 624 | 108 | 18 | 90 | 73.91 | 0.17 |
| APN2/MAT-IGS | 579 | 378 | 201 | 4 | 197 | 149.81 | 0.19 |
| CAL | 648 | 578 | 70 | 1 | 69 | 48.76 | 0.27 |
| GAP2-IGS | 708 | 569 | 135 | 14 | 121 | 79.71 | 0.42 |
| GAPDH | 767 | 734 | 33 | 2 | 31 | 24.51 | 0.33 |
| GS | 665 | 571 | 87 | 24 | 63 | 43.83 | 0.39 |
| TUB2 | 1231 | 1116 | 115 | 10 | 105 | 67.23 | 0.49 |
| ***C. graminicola s. l.*** |  |  |  |  |  |  |  |
| ACT | 269 | 174 | 82 | 52 | 30 | 30.25 | 0.68 |
| CHS-1 | 280 | 237 | 43 | 24 | 19 | 17.06 | 0.55 |
| nrITS | 454 | 375 | 70 | 37 | 33 | 30.38 | 0.23 |
| TUB2 | 500 | 350 | 145 | 89 | 56 | 58.96 | 0.46 |
| ***C. magnum s. l.*** |  |  |  |  |  |  |  |
| ACT | 276 | 260 | 16 | 13 | 3 | 11.16 | 0.40 |
| CHS-1 | 257 | 247 | 10 | 7 | 3 | 11.16 | 0.40 |
| GAPDH | 241 | 215 | 26 | 14 | 12 | 13.35 | 0.42 |
| HIS3 | 399 | 360 | 39 | 28 | 11 | 26 | 0.44 |
| nrITS | 539 | 533 | 6 | 6 | 0 | 4.16 | 0.40 |
| TUB2 | 536 | 500 | 36 | 24 | 12 | 23.15 | 0.52 |
| ***C. orbiculare s. l.*** |  |  |  |  |  |  |  |
| ACT | 226 | 202 | 21 | 5 | 16 | 18.37 | 0.36 |
| CHS-1 | 280 | 268 | 12 | 2 | 10 | 6.16 | 0.65 |
| GAPDH | 243 | 203 | 40 | 8 | 32 | 32.01 | 0.17 |
| GS | 954 | 810 | 134 | 41 | 93 | 69.13 | 0.53 |
| HIS3 | 386 | 334 | 50 | 10 | 40 | 36.51 | 0.13 |
| nrITS | 529 | 514 | 15 | 4 | 11 | 6.38 | 0.52 |
| TUB2 | 467 | 431 | 36 | 9 | 27 | 19.75 | 0.55 |
| ***C. orchidearum s. l.*** |  |  |  |  |  |  |  |
| ACT | 276 | 244 | 29 | 17 | 12 | 7.06 | 0.52 |
| CHS-1 | 265 | 243 | 22 | 6 | 16 | 11.46 | 0.47 |
| GAPDH | 242 | 195 | 47 | 37 | 10 | 7.99 | 0.77 |
| HIS3 | 402 | 343 | 59 | 31 | 28 | 16.48 | 0.67 |
| nrITS | 539 | 521 | 18 | 10 | 8 | 5.81 | 0.57 |
| TUB2 | 540 | 460 | 79 | 51 | 28 | 17.44 | 0.80 |
| ***C. spaethianum s. l.*** |  |  |  |  |  |  |  |
| ACT | 210 | 152 | 56 | 28 | 28 | 19.44 | 0.77 |
| GAPDH | 240 | 132 | 105 | 28 | 77 | 32.53 | 0.47 |
| nrITS | 482 | 470 | 10 | 4 | 6 | 8.66 | 0.73 |
| TUB2 | 443 | 351 | 92 | 44 | 48 | 33.26 | 0.62 |
| ***C. truncatum s. l.*** |  |  |  |  |  |  |  |
| ACT | 215 | 179 | 36 | 6 | 30 | 22.89 | 0.41 |
| GAPDH | 276 | 186 | 90 | 6 | 84 | 56.86 | 0.46 |
| nrITS | 479 | 459 | 20 | 4 | 16 | 10.17 | 0.53 |
| TUB2 | 434 | 353 | 76 | 9 | 67 | 33.14 | 0.53 |

1 - number of base pairs.
2 - PIV, phylogenetic informativeness values on a per net basis
3 - PImax, optimal divergence time. Values are expressed in arbitrary units.

Most markers employed for *Colletotrichum* systematics comprise partial sequences of orthologous protein-coding genes. These markers are composed of introns flanked by long exons that are highly conserved. The GAPDH, GS, HIS3 and TUB2 markers contain long intronic regions and APN2/MAT-IGS and GAP2-IGS present long intergenic regions. This may explain why these markers are more variable than the protein-coding loci. While protein-coding loci may be useful for providing support along the backbone of the phylogeny within a species complex or across the genus, markers with variable introns and intergenic sequences are preferable for application at lower taxonomic levels (Schmitt et al., 2009).

### 3.2 DNA barcoding feasibility

GAPDH had the largest barcode gap distance in seven of the 11 species complexes evaluated. The percentage overlap between intra- and interspecific distances was less than 20%, with the exception of the *C. gigasporum* and the *C. gloeosporioides* species complexes (28.6% and 29.2%, respectively). GS has the highest barcode distance with the lowest overlap for the *C. gigasporum* (0.12 and 12.1%) and *C. orbiculare* (0.08 and 2.7%) complexes. APN2/MAT-IGS had the largest barcode gap distance (0.15) and the smallest overlap percentage (3.26%) within the *C. gloeosporioides* species complex, making it the best candidate barcode locus for the complex HIS3 had the largest barcode gap (0.026) within the *C. orchidearum* complex with a relatively low overlap (16.9%), although CHS-1 had a comparable barcode gap (0.025) with only a slightly higher overlap (21.6%). While other markers, such as HIS3 and TUB2, are good candidate secondary barcode markers in several complexes, nrITS was universally the poorest barcode candidate with the lowest barcode gap distance within all species complexes.

Our results demonstrate that selecting a universal barcode marker for all *Colletotrichum* species complexes among the markers currently being used is not possible. An illustration of this is GAPDH. This marker is the best candidate barcode marker for the majority of complexes, however it is among the worst barcode candidates for the *C. gloeosporioides* species complex. While our results are not in agreement with Cai et al. (2009), which carried out the first study evaluating the markers to discriminate species within *C. gloeosporioides* species complex, only five species were included in their analyses and the Musae and Kahawae clades (*sensu* Weir et al. (2012)) were treated as single species. Moreover, while GAPDH was chosen as the best marker relative to EF1α, ACT, CHS-1 and nrITS, the latter three markers perform very poorly for species delimitation in the *C. gloeosporioides* complex (Vieira et al., 2017) and, therefore, we did not include them in any of our analyses. While GAPDH, with its large barcode gap and small overlap along with the ease with which it can be amplified and sequenced, makes it among the best barcode candidates across *Colletotrichum* as a whole, the selection of the best barcode markers is dependent on the species complex.

The search for a universal barcode locus for the genus will require a comparative analysis across the genomes of several species in all species complexes. Intergenic sequences in syntenic regions of the genome are good candidates if APN2/MAT-IGS and GAP-IGS are any indication. Intergenic sequences may provide fast-evolving phylogenetic markers to be used for population genetic and phylogenetic studies on fungal species complexes (Magain et al., 2017).

### 3.3 Optimal markers for phylogenetic inference

The most informative markers differ among the species complexes within *Colletotrichum*. Net phylogenetic informativeness profiles and their respective ultrametric trees are presented in Fig. 2. PIV and PImax values are summarized in Table 1. The GSI analyses illustrate the ability of a given marker to recover species as monophyletic is dependent on the species complex (Fig. 3, GSI values presented in Supplementary File S2). Most species that were monophyletic in the multilocus tree with all markers (GSI near to 1) were also recovered as monophyletic when only the most informative markers were concatenated. In parallel, the BCAs revealed that the proportion of markers supporting individual species-level clades (expressed as concordance factors) increase when the less informative markers are removed from the analyses.

**Fig. 1.**
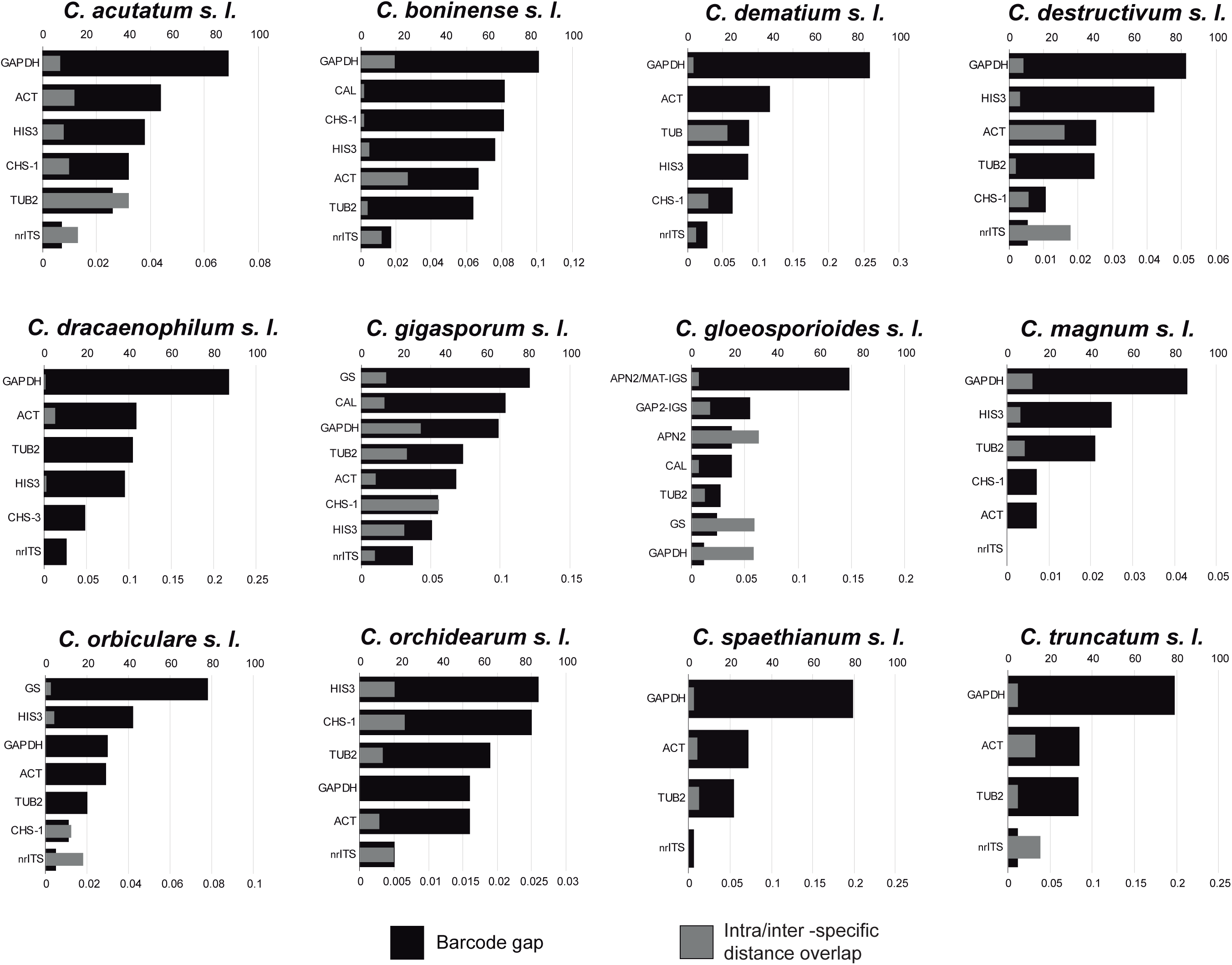
Barcode gap and distance overlap between the intra- and inter-specific distances. Values were calculated based on the intra- and inter-specific distances frequencies distribution of each *Colletotrichum* species complex.

**Fig. 2.**
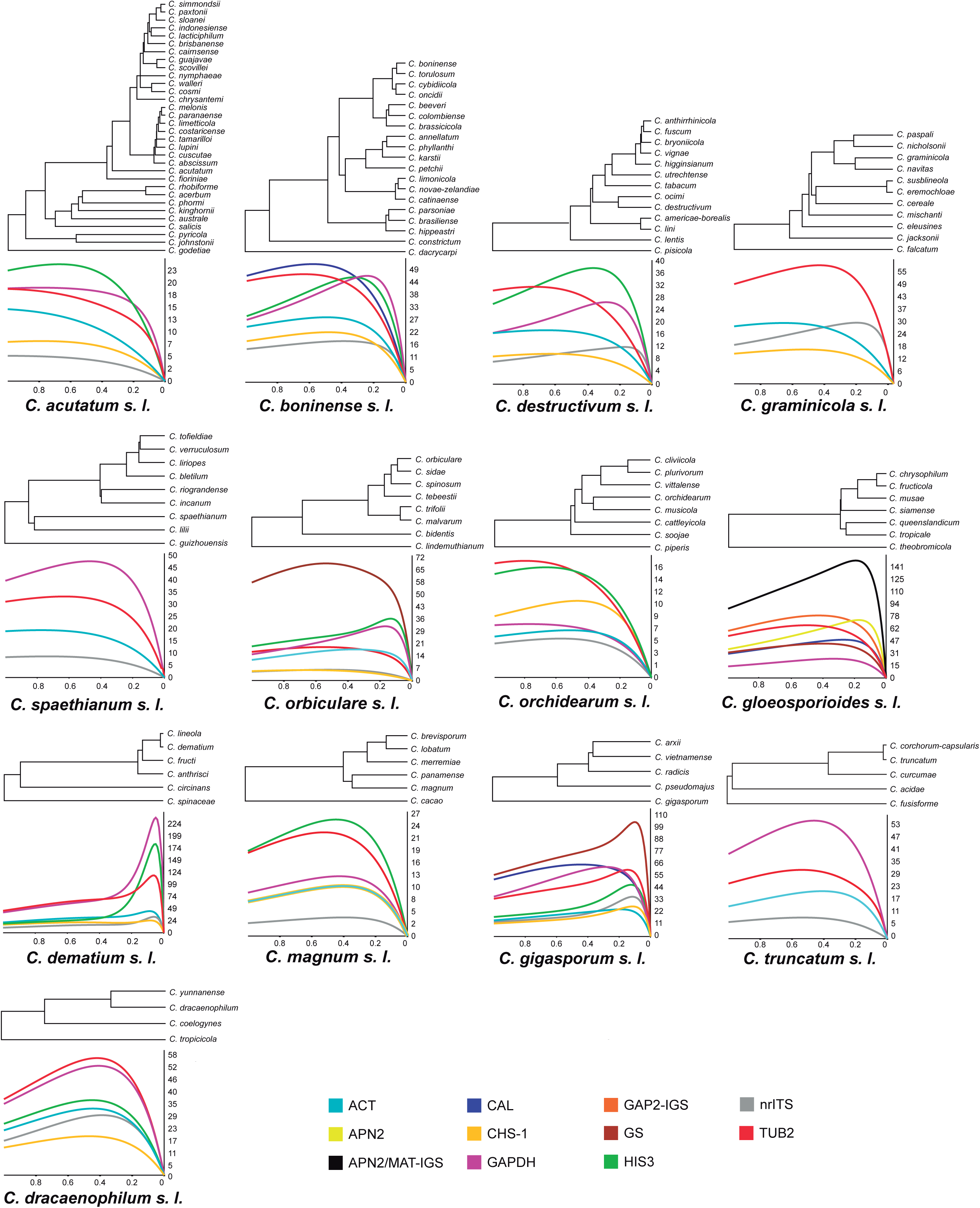
Ultrametric trees and net phylogenetic informativeness profiles of markers used for phylogenetic studies of 13 *Colletotrichum* species complexes. Values on the X-axes correspond to the relative timescale (0—1) based on the root-to-tip distance. Values on the Y-axes represent net phylogenetic informativeness values (10^-6^) in arbitrary units.

**Fig. 3.**
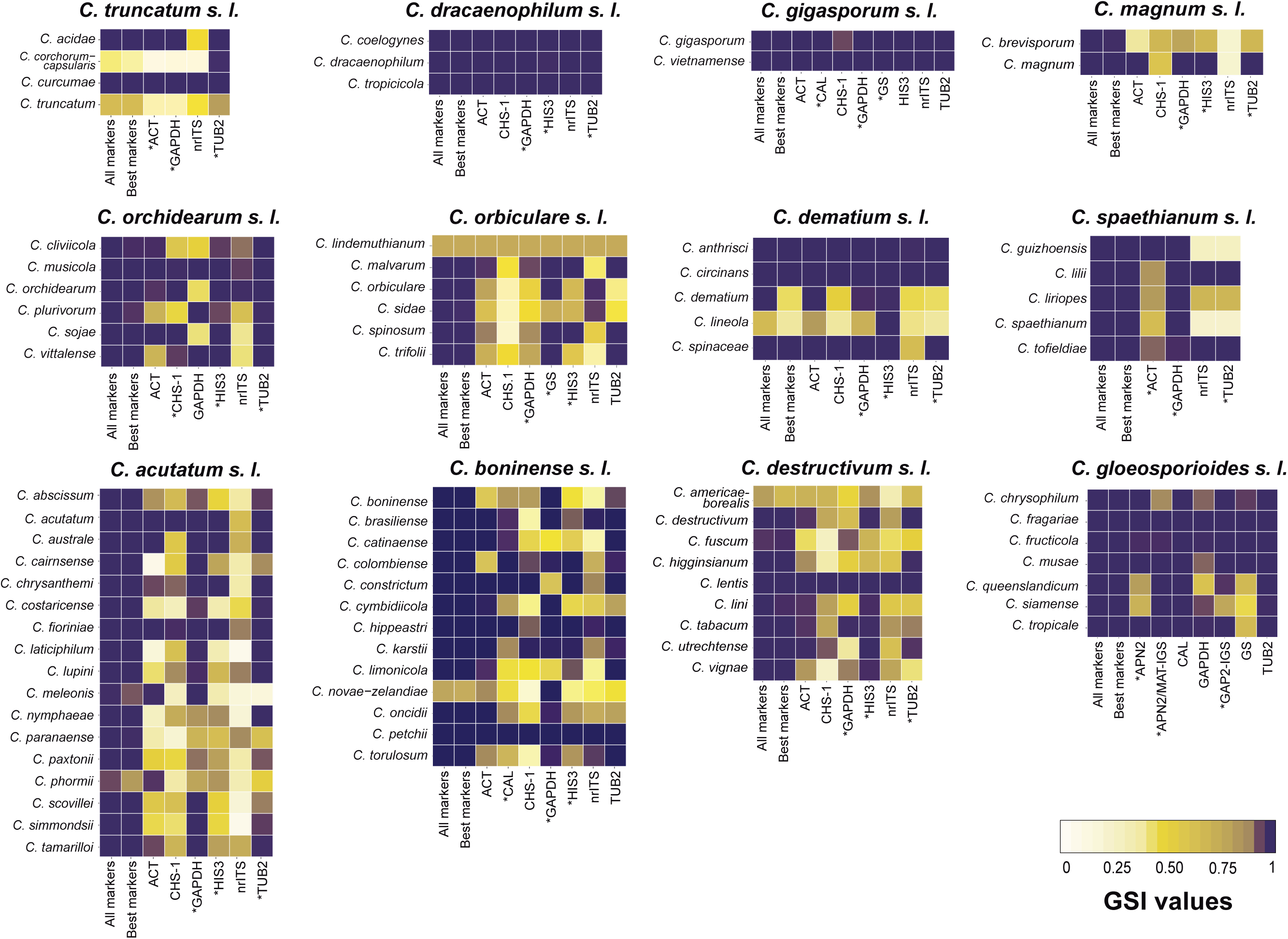
Heat map of the Genealogical Sorting Indices (GSI) by *Colletotrichum* species complex. GSIs of 1000 bootstrap trees were calculated with 100 permutations. Rows correspond to species, and columns correspond to individual markers and concatenated datasets (all markers and best markers). Asterisks represent the best markers for each complex.

Since the suite of markers differ by species complex and the performance of a given marker differs among the species complexes, the results of the phylogenetic informativeness profiling, GSI, Bayesian concordance analyses and selection of the minimum set of markers for robust phylogenetic inference are summarized below for each species complex.

#### 3.3.1 Colletotrichum acutatum s. I

HIS3, GAPDH and TUB2 are the most phylogenetically informative markers within the *C. acutatum* species complex, with PImax at 0.67, 0.78 and 0.99, respectively (Fig. 2). All markers currently used for systematics of the C*. acutatum* complex have an optimum inferential timescale varying from 0.67 to 0.99, which is useful to resolve deeper relationships. However, the absence of markers with lower PImax negatively impacts the our ability to infer relationships among recently diverged species. This is evident by the low internal node support within Clade 1 and Clade 2 *sensu* Damm et al. (2012a), which are the most recently diverged lineages within the complex (Supplementary File S3). Species are strongly supported in the concatenated multilocus analysis (ML support ≥ 70%, Supplementary File S3), whereas the relationship among them are poorly resolved even when only the best markers were used to build the tree. In contrast, nodes throughout Clade 5 are strongly supported. The inclusion of a marker with PImax near to 0.2 could improve the support on the deeper nodes of the Clades 1 and 2.

Most species in *C. acutatum s. l.* could be resolved by the majority of the markers only when the optimal markers are combined (Fig. 4B). Seven (*C. austral*, *C. chrysanthemi*, *C. fioriniae*, *C. johnstonii*, *C. lupini*, *C. nymphaeae* and *C. tamarilloi*) among the 17 multiple-isolate species were supported by the majority of individual genes (CF≥0.66) in the BCA when all six loci were used in the analysis (Fig. 4A), which means that four out of six markers support the monophyly of these species. The CF increases when the analyses included only HIS3, GAPDH and TUB2, and only four (*C. cosmi*, *C. costaricense*, *C. paranaense* and *C. phormii*) out of 17 multiple-isolate species presented CF<0.66 (Fig. 4B). The GSI (Fig. 3) indicates that *C. melonis* is recovered by GAPDH and *C. phormii* by HIS3. The GSI for *Colletotrichum costaricense* was nearly 1 for both GAPDH and TUB2 (0.95 and 0.99, respectively) and less than 0.5 for all other markers. This is consistent with a CF of 0.63, indicating that nearly 2 of the 3 loci included in the analysis support the monophyly of the species. *Colletotrichum paranaense* had a low GSI for most markers and CF=0.22, which means that no individual marker fully supports this species as monophyletic. However, *C. paranaense* is strongly supported in the multilocus concatenated analysis and has a high GSI when analyses included all or the best markers. These results clearly show that the multilocus trees masks the incongruences among the individual gene trees or the use of markers with low phylogenetic signal.

**Fig. 4.**
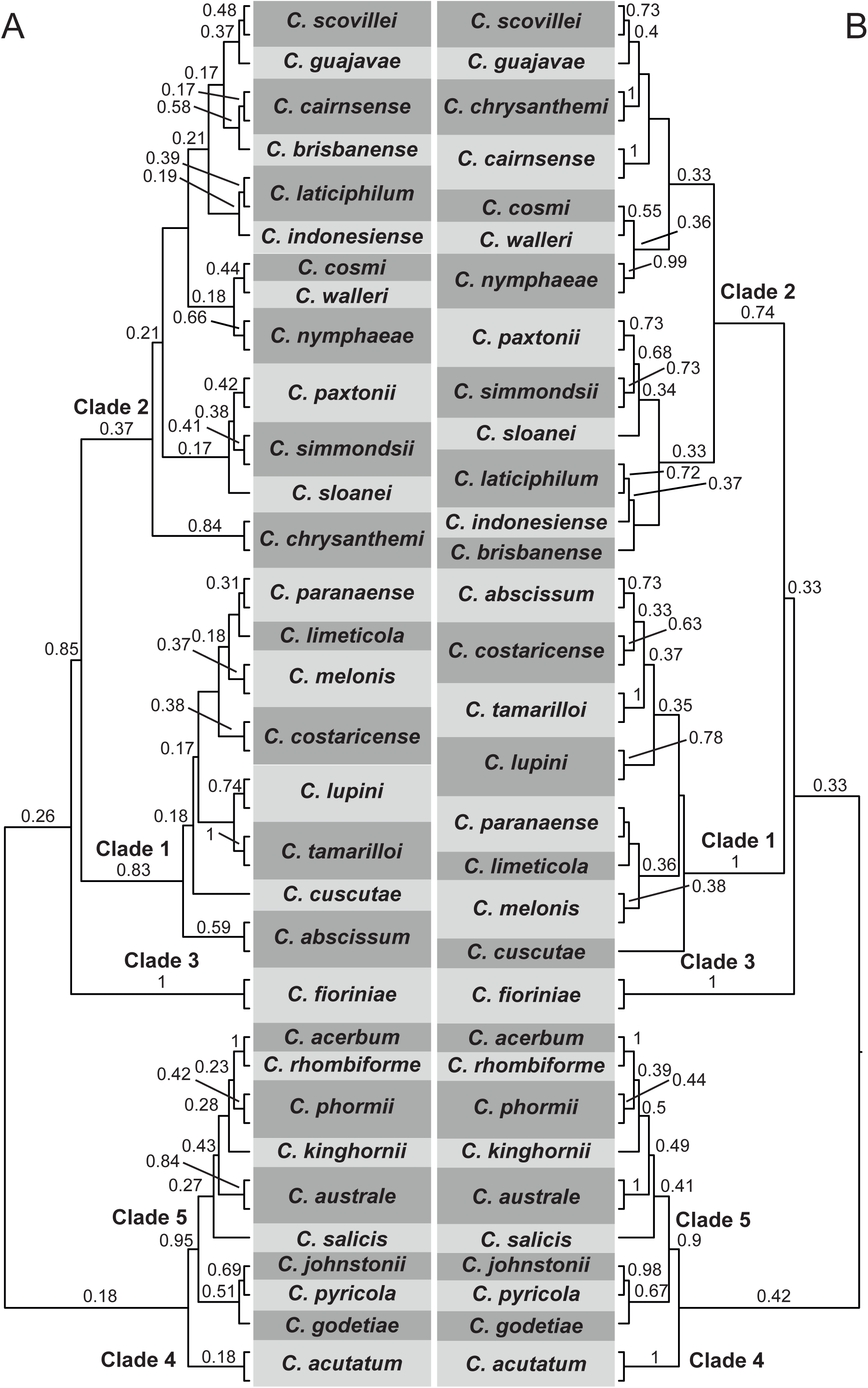
Primary concordance trees resulting from the Bayesian concordance analyses including isolates from the *C. acutatum* complex. A. All markers (ACT, CHS-1, GAPDH, HIS3, ITS and TUB2). B. Best markers (GAPDH, HIS3 and TUB2). Concordance factors are shown above the branches that were resolved by at least one marker (≥0.16 for all markers and ≥0.33 for the best markers).

*Colletotrichum paranaense* was described by Bragança et al. (2016) as a species closely related to *C. limetiicola* and *C. melonis* (Damm et al., 2012a). Phylogenetic species recognition was not employed in the delimitation of *C. paranaense*. The species was recognized based on the topology of the multilocus tree and differed from *C. limetticola* and *C. melonis* by percentage identity according to blastn searches. Our results suggest that the relationship among these species and the delimitation of *C. paranaense* needs to be revisited.

#### 3.3.2 Colletotrichum boninense s. I

CAL, TUB2 and GAPDH were the most informative markers to resolve species within *C. boninense s. l.* (Fig. 2). All three markers are comparable in levels of phylogenetic informativeness, with PIV ranging from 47 to 53. However, these markers differ with respect to the optimal timescale at which they are informative. CAL and TUB2 peaked at 0.58 and 0.63, respectively, providing more signal towards the root of the phylogeny. In contrast, GAPDH reached maximum informativeness at 0.24, which is more useful for resolving species in this complex, since the divergence epoch for most species is near 0.2.

Most species can be resolved by most of the markers according to the concordance analyses that include all markers and only the best markers (CF>0.57 and >0.66, respectively) (Fig. 5B). GAPDH supports most species as monophyletic (GSI 0.98—1) (Fig. 2), with the exception of three species (*C. catinaense*, *C. constrictum* and *C. limonicola*). According to Damm et al. (2012b), all the species within the *C. boninense* species complex can be identified by sequencing GAPDH. In contrast, our results show that *C. catinaenese*, *C. constrictum* and *C. limonicola* are weakly supported as monophyletic by GAPDH (GSI = 0.51, 0.69 and 0.59, respectively) and could only be recovered as monophyletic in analyses of TUB2, HIS or ACT. *Colletotrichum cymbidiicola* and *C. novae-zelandiae* were recovered as monophyletic with GAPDH and *C. limonicola* by ACT and TUB2 (Fig. 3). Thus, concordance factors for these species remained low even when the best markers were used in the BCA. Additionally, *C. novae-zelandiae* was the only species for which GSI<1 in both multilocus analyses (Fig. 3). *Colletotrichum catinaense* could be resolved by two of the best markers (CAL and TUB) and by ACT, which led the CF to shift from 0.56 in the BCA with all markers to 0.73 when only the best markers were included (Fig. 5). The use of CAL, TUB2 and GAPDH leads to a slight increase in support across the multilocus ML tree when compared with the tree reconstructed with the whole dataset (Supplementary File S4), with the exception of *Colletotrichum novae-zelandiae*, which remained poorly supported.

**Fig. 5.**
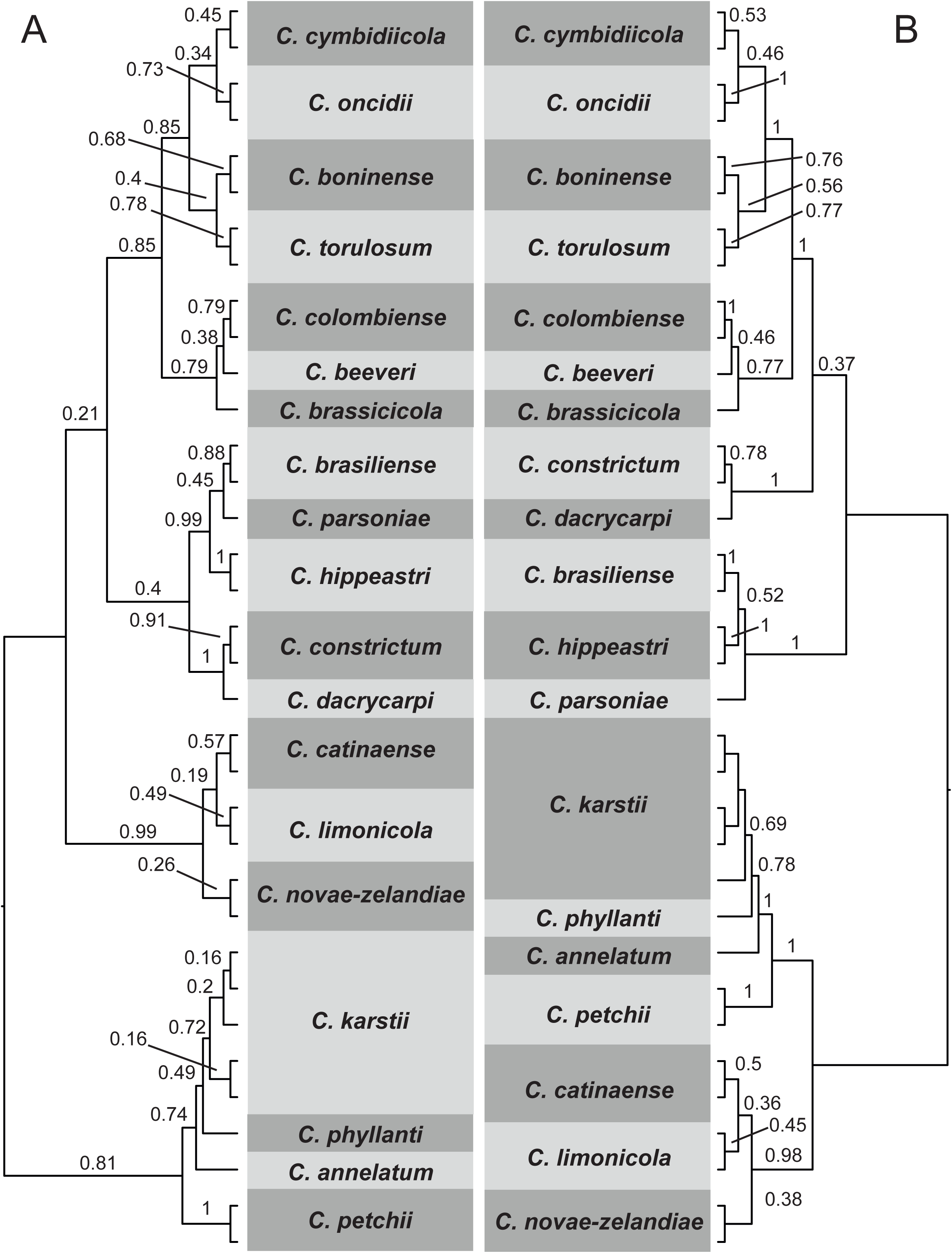
Primary concordance trees resulting from the Bayesian concordance analyses including isolates from *C. boninense* complex. A. All markers (ACT, CAL, CHS-1, GAPDH, HIS3, ITS and TUB2). B. Best markers (CAL, GAPDH and TUB2). Concordance factors are shown above the branches that were resolved by at least one marker (≥0.14 for all markers and ≥0.33 for the best markers).

#### 3.3.3 Colletotrichum dematium s. I

GAPDH, HIS3 and TUB2 presented high and sharp profiles (Fig. 2), with PImax 0.05—0.06 and were the most useful markers to discriminate species within the *C. dematium* species complex. HIS3 was the only marker able to discriminate all species in the complex. nrITS, CHS-1 and ACT presented low and flat curves and were the least informative markers (Fig. 2). *Colletotrichum anthrisci* and *C. circinans* could be resolved by all markers (CF=1, Fig. 6), while *Colletotrichum spinaceae* was resolved as monophyletic by all markers except nrITS. The exclusion of nrITS in the BCA increased the CF from 0.86 to 1. The relationship between *Colletotrichum dematium* and *C. lineola* was not clearly resolved in the multilocus analyses. Although *C. dematium* isolates were well supported as monophyletic in the tree inferred from all markers, *C. lineola* isolates remained paraphyletic (Supplementary File S5). When only GAPDH, HIS3 and TUB2 were considered, the isolates of *C. dematium* and *C. lineola* were placed together in a poorly supported clade. *Colletotrichum lineola* was recovered as monophyletic only by HIS3 (GSI=1). Both ACT and GAPDH could also resolve this species, albeit with low GSI (0.83 and 0.73 respectively). In contrast, these three markers recover *C. dematium* with high level of monophyly (GSI 0.97—1). *Colletotrichum lineola* presented low monophyly level in all multilocus GSI analysis and *C. dematium* was not monophyletic when only the best markers were considered.

**Fig. 6.**
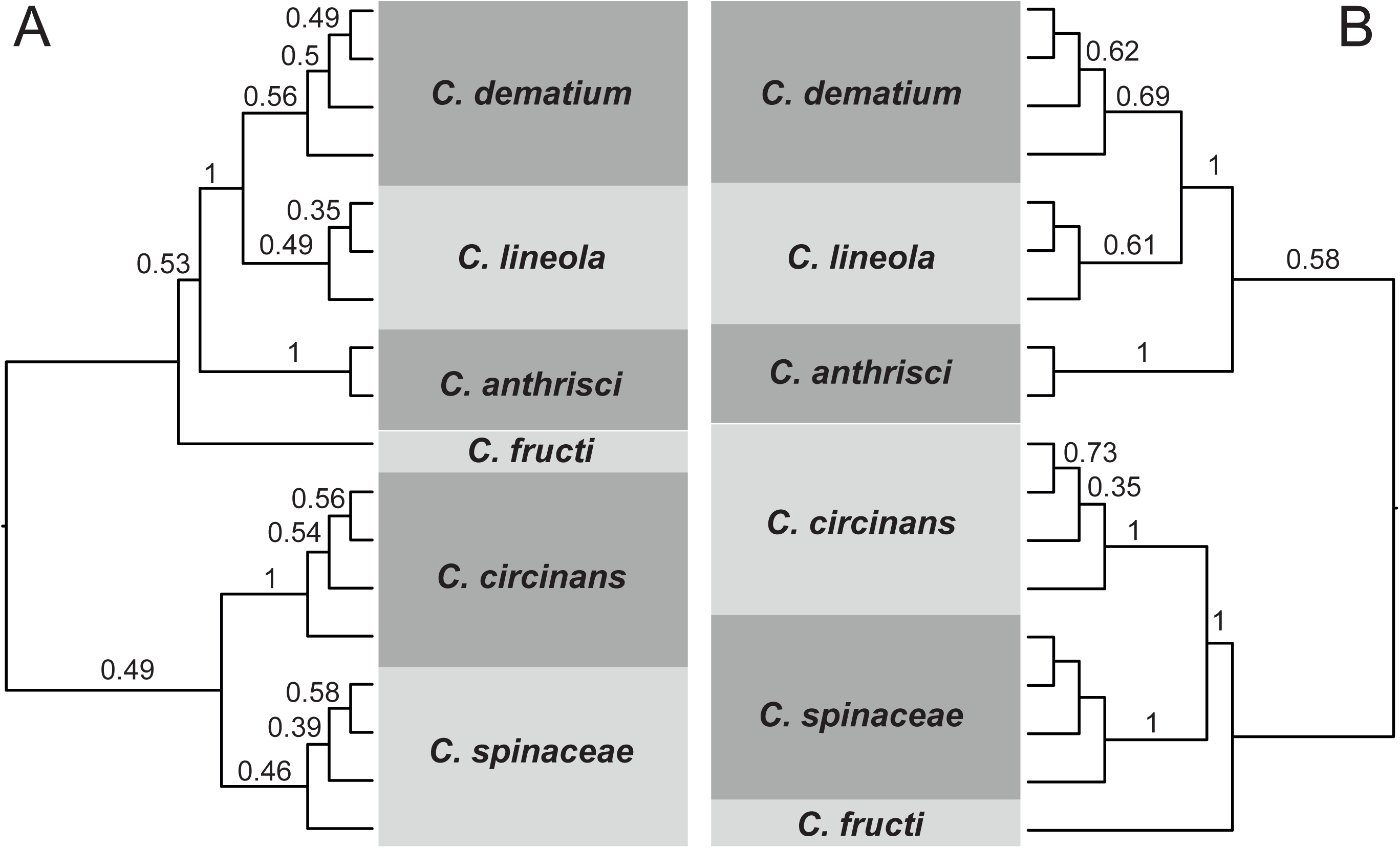
Primary concordance trees resulting from the Bayesian concordance analyses including isolates from the *C. dematium* complex. A. All markers (ACT, CHS-1, GAPDH, HIS3, ITS and TUB2). B. Best markers (GAPDH, HIS3 and TUB2). Concordance factors are shown above the branches that were resolved by at least one marker (≥0.16 for all markers and ≥0.33 for the best markers).

In the Damm et al. (2009) study, the isolates of *C. dematium* and *C. lineola* are separated into two short-branch clades. They chose to retain both taxa, but raised the hypothesis that *C. dematium* and *C. lineola* are different populations within the same species. We performed the GSI analyses with the multilocus trees combining *C. dematium* and *C. lineola* in the same group (data not shown). The group was recovered as monophyletic when all markers and only the best markers are concatenated (GSI=1 and 0.99 respectively), which support the hypothesis that *C. dematium* and *C. lineola* are the same species. *Colletotrichum eryngiicola*, *C. hemerocallidis*, *C. insertae*, *C. quiquefoliae* and *C. sonchicola* were placed together with *C. dematium* and *C. lineola* in a polytomous clade in Samarakoon et al. (2018) study, which indicates that those species may also be members of the group *C. dematium*/ *C. lineola*. These species were not included in our analyses due to absence of HIS3 and TUB2 sequences. In the future, species boundaries within this lineage need to be revisited.

#### 3.3.4 Colletotrichum destructivum s. I

HIS3, TUB2 and GAPDH were the most informative markers to resolve species within *C. destructivum* complex (Fig. 2). GAPDH and HIS3 possess the phylogenetic signal to resolve shallow clades (PImax=0.28 and 0.37 respectively), whereas TUB2 performs better in the deep branches (PImax=0.73). HIS3 was the most informative marker and was able to six (*C. destructivum*, *C. lentis*, *C. lini*, *C. tabacum*, *C. utrechtense* and *C. vignae*) out of the nine species analyzed as monophyletic (GSI 0.98—1) (Fig. 3). Although GAPDH is one of the best markers, only four species (*C. fuscum*, *C. lentis*, *C. tabacum* and *C. vignae*) were highly monophyletic in the topologies provided by this locus (GSI 0.9—1) (Fig. 3). *Colletotrichum americae-borealis* was not monophyletic in the topology of any single or multi-locus trees (GSI 0.34—0.89). Four (*C. destructivum*, *C. lentis*, *C. tabacum*, *C. utrechtense*) out of nine multiple-isolates species could be recovered by most of the genes in the concordance tree (CF≥0.66) inferred from all markers (Fig. 7A). These same four species plus *C. vignae* were recovered by most of the genes (CF≥0.66) when only the optimal markers were considered. Moreover, the CF and the bootstrap supports of the internal nodes increased when only the best markers were combined (Fig. 7 and Supplementary File S6).

**Fig. 7.**
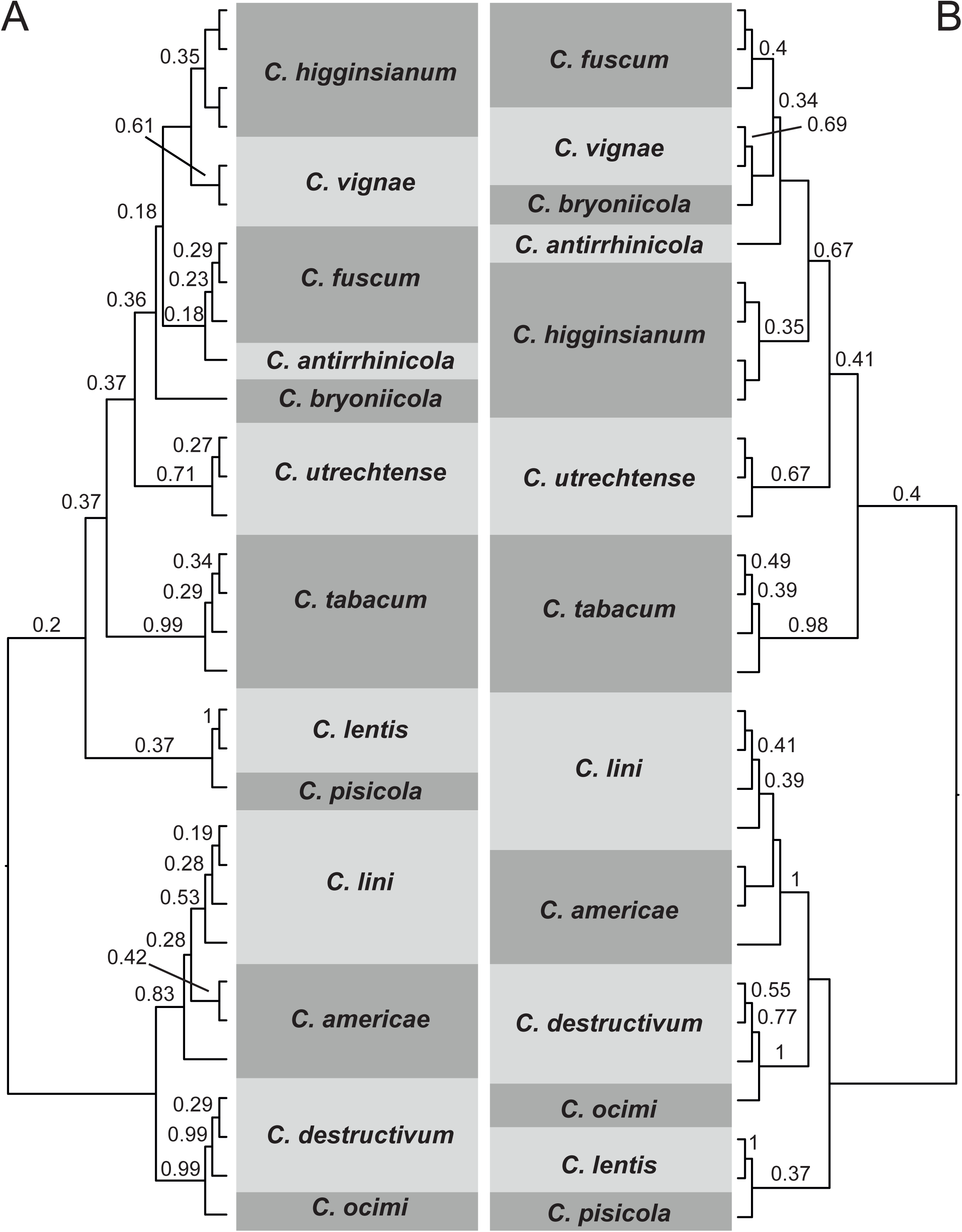
Primary concordance trees resulting from the Bayesian concordance analyses including isolates from the *C. destructivum* complex. A. All markers (ACT, CHS-1, GAPDH, HIS3, ITS and TUB2). B. Best markers (GAPDH, HIS3 and TUB2). Concordance factors are shown above the branches that were resolved by at least one marker (≥0.16 for all markers and ≥0.33 for the best markers).

The clade comprised by *C. americae-borealis* and *C. lini* forms a polytomy when only the three best markers were used (Supplementary File S6). We also tested the inclusion of ACT or nrITS in the concatenated dataset. However, this clade remains a polytomy and can only be resolved when all markers are concatenated. The overall GSI values also reduced when ACT or nrITS was included in the multilocus analysis (data not shown). Thus, it is not reasonable to include ACT and nrITS in the analysis due to their low phylogenetic informativeness. Other markers with greater phylogenetic signal need to be tested for this complex in order to resolve the relationships among the unresolved clades and clarify the species identities.

#### 3.3.5 Colletotrichum dracaenophilum s. I

TUB2, GAPDH and HIS3 presented the highest PIV (58, 52 and 36 respectively) and were the most informative markers to distinguish species within the *C. dracaenophilum* complex (Fig. 2). All the markers peaked around the same timescale (PImax 0.39—0.46) and provide robust support for relationships in both deep and shallow nodes. Although TUB2, GAPDH and HIS3 were the most informative markers, all the markers are informative enough to discriminate species in *C. dracaenophilum s. l.* (GSI 0.99—1) (Fig. 3). This is corroborated by the high concordance factors (CF=1) in BCAs (Fig. 8) and 100% support in multilocus analyses (Supplementary File S7) when all or the best markers were combined.

**Fig. 8.**
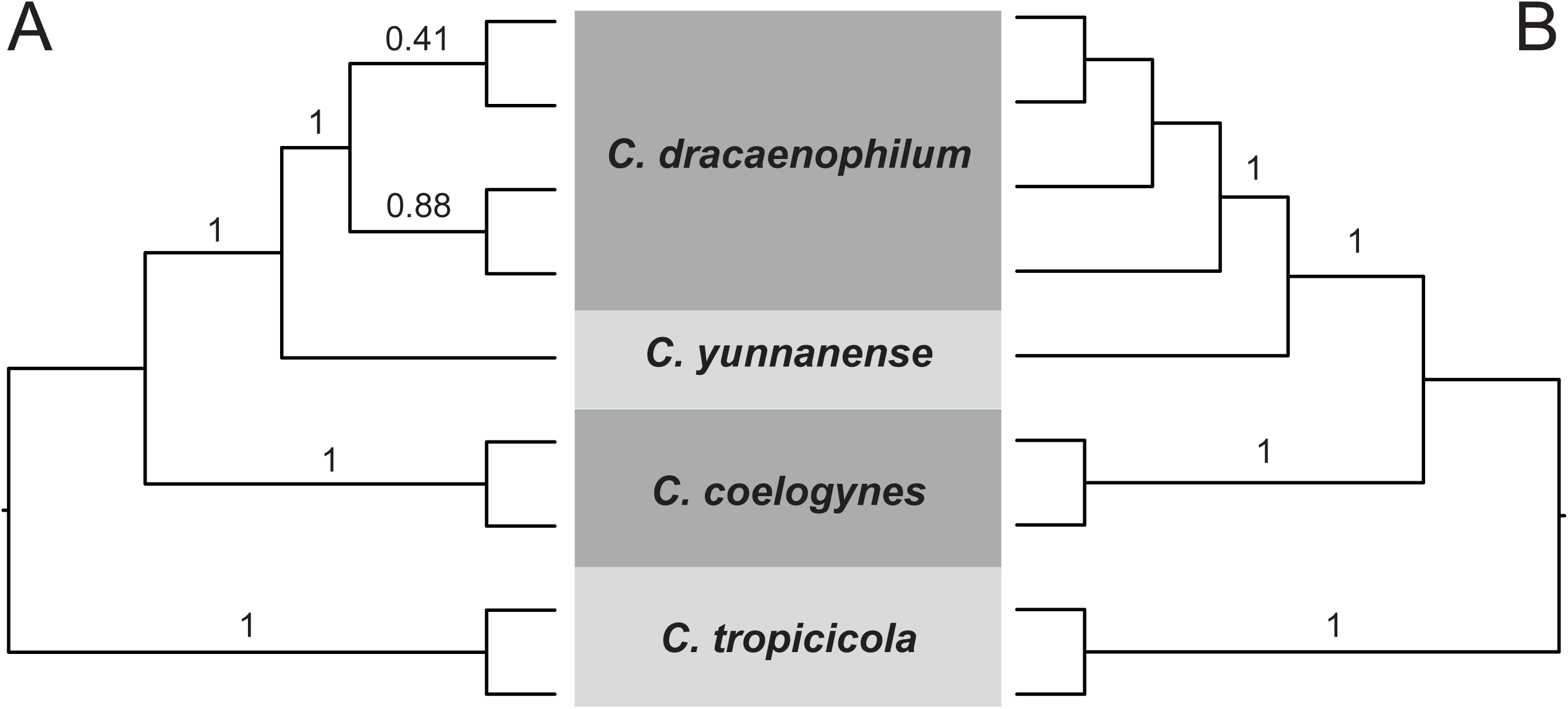
Primary concordance trees resulting from the Bayesian concordance analyses including isolates from the *C. dracaenophilum* complex. A. All markers (ACT, CHS-1, GAPDH, HIS3, ITS and TUB2). B. Best markers (GAPDH, HIS3 and TUB2). Concordance factors are shown above the branches that were resolved by at least one marker (≥0.16 for all markers and ≥0.33 for the best markers).

#### 3.3.6 Colletotrichum gigasporum s. I

GS, CAL and GAPDH were the most powerful markers to discriminate species within *C. gigasporum s. l.* (Fig. 2), with PImax at 0.1, 0.44 and 0.28 respectively. Although GS, CAL and GAPDH were the most informative markers, all the markers and the concatenated datasets were able to discriminate all species within the *C. gigasporum* complex (GSI 0.99—1) (Fig. 3), as reported by Liu et al. (2014). All species presented maximum CF (Fig. 9) and ML support (Supplmentary File S8) independently of which set of markers was included in these analyses.

**Fig. 9.**
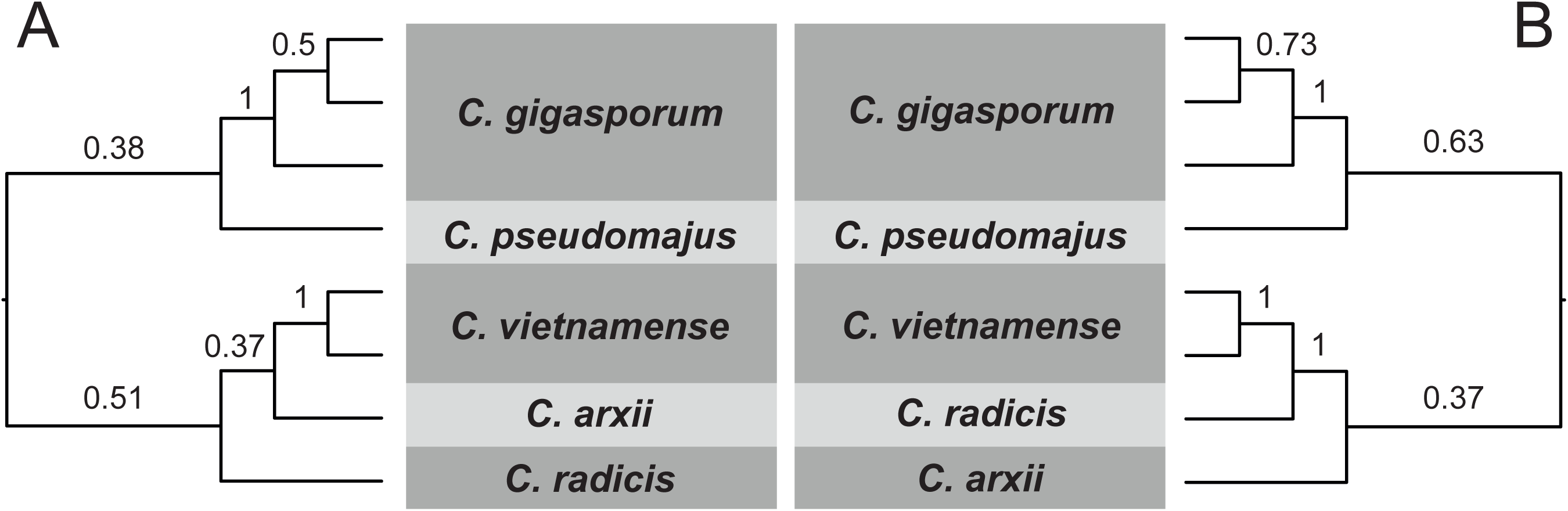
Primary concordance trees resulting from the Bayesian concordance analyses including isolates from the *C. gigasporum* complex. A. All markers (ACT, CAL, CHS-1, GAPDH, GS, HIS3, ITS and TUB2). B. Best markers (CAL, GAPDH and GS). Concordance factors are shown above the branches that were resolved by at least one marker (≥0.13 for all markers and ≥0.33 for the best markers).

#### 3.3.7 Colletotrichum gloeosporioides s. I

APN2/MAT-IGS, GAP2-IGS and APN2 were the most informative markers to separate species within the *C. gloeosporioides* species complex (Fig. 2). The informativeness profiles indicate APN2/MAT-IGS and APN2 are of peak informativeness at 0.19 and 0.17, respectively, and are informative for shallow divergences, whereas GAP2-IGS has an optimal timescale at 0.42 and provides more signal for resolving deep nodes. This range in values for peak informativeness led to high support at both deep and shallow nodes when only the most informative markers were used for phylogenetic inference (Supplementaray File S9). APN2/MAT-IGS could separate all the species included in the analyses, although only *C. fragariae*, *C. queenslandicum*, *C. siamense* and *C. tropicale* reached a GSI of 1 (Fig. 3). Moreover, all species were recovered as monophyletic in all multilocus analyses. *Colletotrichum siamense* was the only species that could not be recovered by most of the markers, and its CF reduced from 0.59 in all-markers analysis to 0.38 in the best-markers analysis (Fig. 10B). The CF reduced due to removing CAL and TUB2, which also recover *C. siamense* as monophyletic (GSI=0.99).

**Fig. 10.**
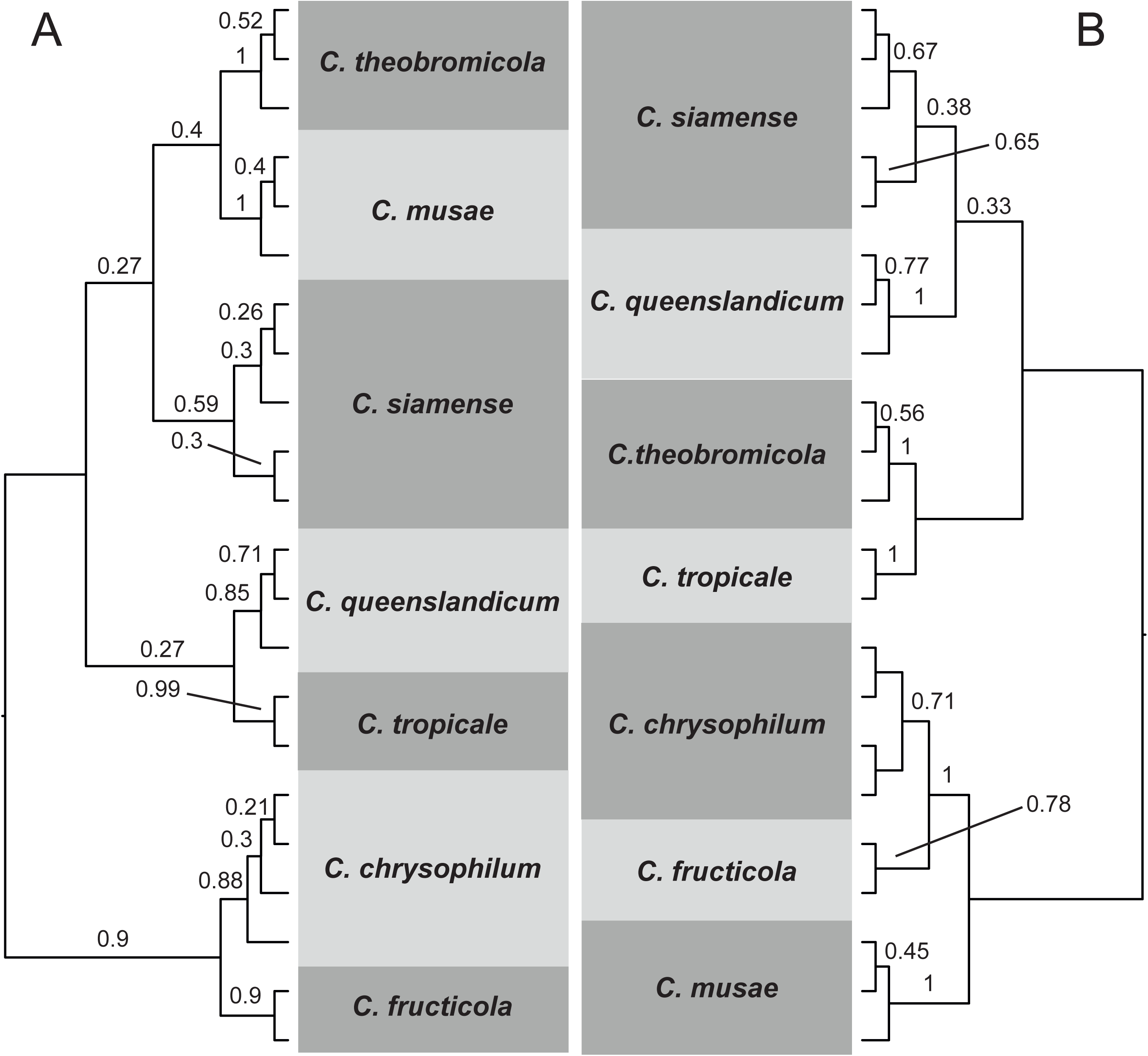
Primary concordance trees resulting from the Bayesian concordance analyses including isolates from the *C. gloeosporioides* complex. A. All markers (APN2, APN2/MAT-IGS, CAL, GAPDH, GAP2-IGS, GS and TUB2). B. Best markers (APN2, APN2/MAT-IGS and GAP2-IGS). Concordance factors are shown above the branches that were resolved by at least one marker (≥0.14 for all markers and ≥0.33 for the best markers).

ACT, CHS-1 and nrITS were not included in our analyses because these markers were previously reported as the worst markers to distinguish species in *C. gloeosporioides* complex (Vieira et al., 2017). In the present study, rather than restating the results of Vieira et al. (2017), we evaluated if the seven markers proposed by those authors are needed for diversity studies and species assignment. We determined sequences of APN2/MAT-IGS, GAP2-IGS and APN2 are sufficient to resolve the species in *C. gloeosporioides s. l.*. However, sequences of GAP2-IGS and APN2 are not available for all species within *C. gloeosporioides s.l..* These data should be generated for all species within the species complex.

*Colletotrichum gloeosporioides s. l.* is the most studied species complex in the genus. More than 10 different markers have been used for systematics and taxonomy of the *C. gloeosporioides* complex over the last 10 years. However, there is no agreement about which markers should be used for species recognition. The first attempt to determine the best markers was done by Cai et al. (2009), in which GAPDH was the best marker to discriminate species in *C. gloeosporioides s. l.*, and the set composed by ACT, GAPDH, nrITS and TUB2 was recommended for multilocus analysis. Weir et al. (2012) highlighted that although GAPDH is one of the most effective markers to distinguish species within *C. gloeosporioides* complex, the combination with GS is necessary to distinguish some species. Our study shows that GAPDH was the least informative marker among the ones included in this study for *C. gloeosporioides s. l.*. This marker was the least variable (Table 1), and had the smallest barcode gap and the largest overlap distance (Fig. 1). *Colletotrichum fragariae* was the only species recovered as monophyletic (GSI=1) by this marker (Fig 2). Based on this, GAPDH is considered one of the worst barcode candidates for the *C. gloeosporioides* complex among the markers tested in the present study.

More recently, several studies demonstrate the singular ability of APN2/MAT-IGS to discriminate species in the *C. gloeosporioides* complex (Sharma et al., 2013, 2015; Vieira et al., 2014), although others previously demonstrated the utility of this marker (Du et al., 2005; Rojas et al., 2012; Silva et al., 2012). The main issue in using APN2/MAT-IGS was the splitting out of *C. siamense* in a species complex, in which several monophyletic lineages could be revealed by the phylogeny inferred by this marker (Sharma et al., 2013). Although the multilocus analysis had been done, the lineages identity was confirmed only based on the APN2/MAT-IGS, since this marker performs good as well as the multilocus matrix. Other studies use the same criteria to discriminate species and several species within *C. siamense s. l.* were described (Sharma et al., 2015; Vieira et al., 2014). Later, Liu et al. (2016) use coalescent methods for phylogenetic species delimitation and synonymize all those species in the complex into *C. siamense*. It was revealed incongruences among the APN2/MAT-IGS tree and other individual gene trees. The study clarifies how multilocus analyses can mask discordances among individual gene trees and lead to species misidentification.

The combination of APN2/MAT-IGS and GS were proposed as the barcode to delimit species within *C. gloeosporioides* complex (Liu et al., 2015). These two markers were collectively powerful enough to discriminate the 22 species included in the study and produced the same topology as that inferred from a 6 marker multilocus dataset (ACT, CAL, GAPDH, GS, nrITS and TUB2). We tested the combination of APN2/MAT-IGS with each remaining marker individually and all resulting trees were similar in topology (data not shown). Some species, such as *C. siamense*, are polyphyletic in our GAPDH and GS trees, which belies the incongruence with the multilocus and the other individual gene trees as presented by Silva et al. (2012) and Vieira et al. (2017). Mating-type associated markers, such as APN2/MAT-IGS and MAT1-2, had fast evolutionary rates and high variability, and may dominate the topology of multilocus trees (Liu et al., 2016). Since the combination of APN2/MAT-IGS with any other marker produces similar topologies, the combination with any other marker besides GAPDH and GS is preferable in order to avoid inconsistencies in species delimitation within the *C. gloeosporioides* complex. While we find that APN2 is another informative marker that should be incorporated into diversity studies of the *C. gloeosporioides* complex, the proximity of this marker to the APN2/MAT-IGS region suggests it is part of the same linkage group and thus may not represent an independent sample of an organisms evolutionary history. If this is the case, linking substitution models and inferred trees across these two loci would be required and the addition of another locus may be necessary to fulfill the expectations under GCPSR.

Although the majority of *Colletotrichum* species belong to the *C. gloeosporioides* complex, identification and description of taxa within this complex is the most problematic. Sequences of the best markers, mainly APN2 and GAP2-IGS, have not been sequenced for the majority of species, which prohibits the detection of these species using these markers. A priority should be to generate sequences of the best markers for all species to avoid further misidentification and the introduction of dubious taxa. Until this happens, the description of new species within the *C. gloeosporioides* complex will likely require sequencing multiple additional markers to be certain of their novelty.

#### 3.3.8 Colletotrichum graminicola s. I

TUB2 was the most informative marker, followed by nrITS and ACT (Fig. 2). In contrast, CHS-1 was ranked as the worst marker for the *C. graminicola* species complex. GSI and BCA analyses could not be performed for this complex because only one isolate per species could be included in the analyses. Other markers such as APN2, APN2/MAT-IGS, GAPDH, HIS3, MAT1-2 and SOD2 were also used to identify species within *C. graminicola* complex (Cannon et al., 2012; Crouch et al. 2009b, c; Crouch and Tomaso-Peterson, 2012; Du et al., 2005; Moriwaki and Tsukiboshi, 2009; O’Connell et al., 2012; Tao et al., 2013). However, sequences for these markers are missing for several species. According to our results for other complexes, APN2, APN2/MAT-IGS, GAPDH and HIS3 are likely to be more powerful markers than those that we could include in the analyses for the *C. graminicola* complex. Sequences for these markers need to be generated for isolates of several species in order to establish a better set of optimal markers for phylogenetic inference and species discrimination in the *C. graminicola* complex.

#### 3.3.9 Colletotrichum magnum s. I

HIS3, TUB2 and GAPDH were the most informative markers to discriminate species within the *C. magnum* complex (Fig. 2). Although HIS3 presented the highest PIV (26), TUB2 presented a higher PImax (0.52 versus 0.44) and is able to separate more species than HIS3. Only *C. brevisporum* and *C. magnum* were accounted for by more than one isolate and could be analyzed in the BCA and GSI analyses (Fig. 3). *Colletotrichum brevisporum* reach high levels of monophyly only in analyses of the concatenated dataset (GSI 0.98—1, Fig. 3) and only one gene can fully resolve this clade (CF=0.4, Fig 11). In contrast, *C. magnum* is recovered as monophyletic by the multilocus dataset and by the best markers. However, the CF reached 1 only when the best markers were concatenated (Fig 11). *Colletotrichum magnum* remained highly supported when only the best markers were used to reconstruct the phylogeny of the *C. magnum* complex (Supplementary File S10B) with only a slight decrease in support at some internal nodes. The species *C. liaoningense* was not included in the analyses due to sequence deposition errors detected by Damm et al. (2019) from the study were *C. liaoningense* was described (Diao et al., 2017). Damm et al. (2019) concluded that *C. liaoningense* needs to be revisited.

**Fig. 11.**
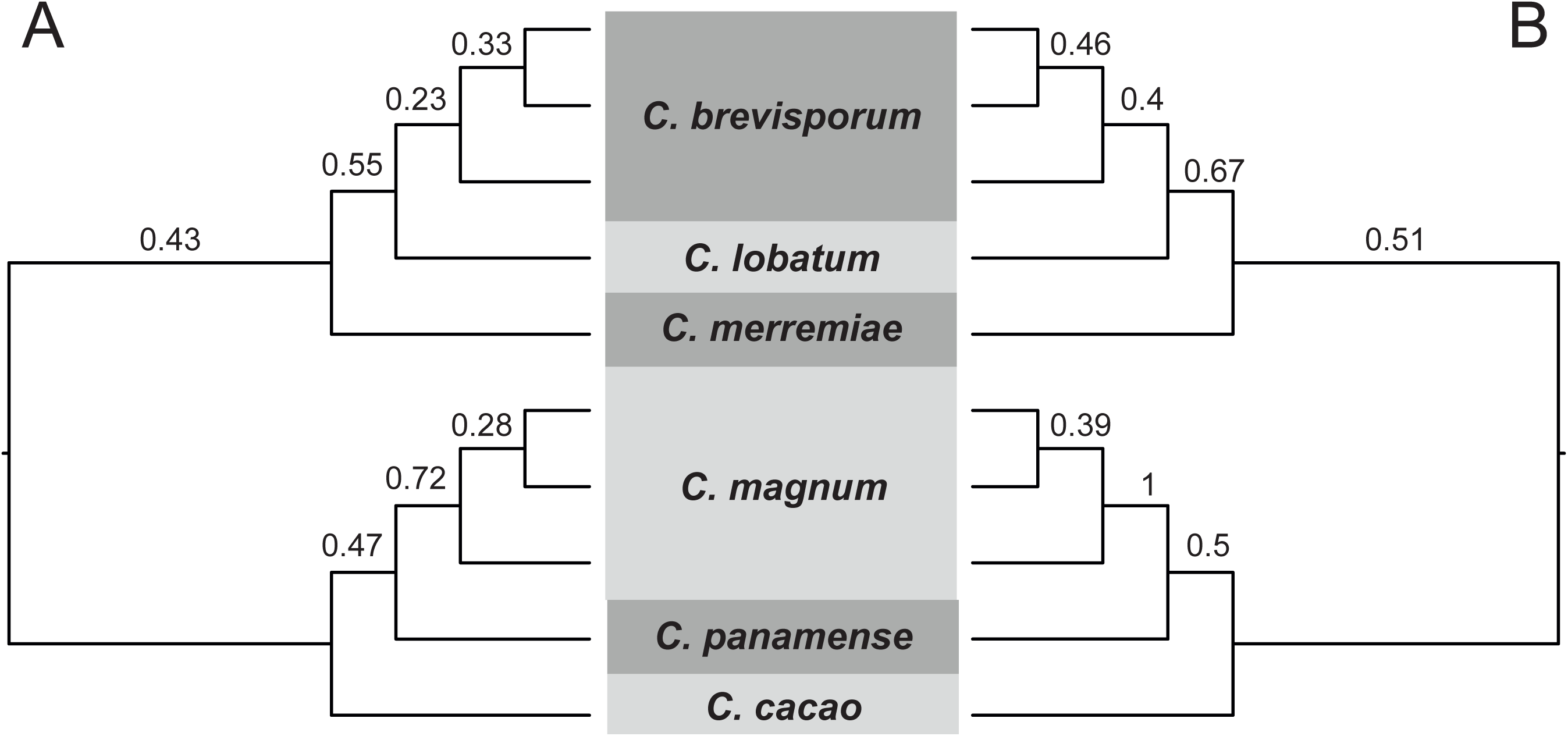
Primary concordance trees resulting from the Bayesian concordance analyses including isolates from the *C. magnum* complex. A. All markers (ACT, CHS-1, GAPDH, HIS3, ITS and TUB2). B. Best markers (GAPDH, HIS3 and TUB2). Concordance factors are shown above the branches that were resolved by at least one marker (≥0.16 for all markers and ≥0.33 for the best markers).

#### 3.3.10 Colletotrichum orbiculare s. I

GS stood out from all markers as the most informative (PIV=69) in the *C. orbiculare* complex, followed by HIS3 and GAPDH (PIV=37 and 32 respectively) (Fig. 2). GS peaks at 0.53 and can discriminate the majority of species and provide robust support for the relationships among them, since the divergence time for most species is about 0.3. On the other hand, HIS3 and GAPDH are useful to discriminate some recently diverged species (PImax=0.13 and 0.17 respectively) such as *C. orbiculare* × *C. sidae* and *C. trifolii × C. malvarum*. GS was the marker that recovered more species as monophyletic (GSI=1), with the exception of *C. sidae* (GSI=0.89) (Fig. 3). *Colletotrichum lindemuthianum* was the only species that was not recovered as monophyletic when all or the best markers were concatenated (both GSI=0.78), which is likely due to the variability within this species as currently circumscribed. Damm et al. (2013) split *C. lindemutianum* into two different lineages (*C. lindemuthianum* 1 and 2), which are observed in the nrITS and GS trees. In our multilocus trees, the isolates CBS133.57 and CBS131.57 where moved to the clade *C. lindemuthianum* 1 and CBS150.28 to the clade *C. lindemuthianum* 2. This result is discordant with that in Damm et al. (2013) and we conclude that the terminal clades within *C. lindemuthianum* represent intraspecific variability. Thus, both *C. lindemuthianum* lineages were not considered separate species in our analysis. The CF of both deep and shallow nodes increases significantly when only the best markers were analyzed (Fig.12). Reducing the set of markers does not cause significant differences in the tree topology or clade support (Supplementary file S10).

**Fig. 12.**
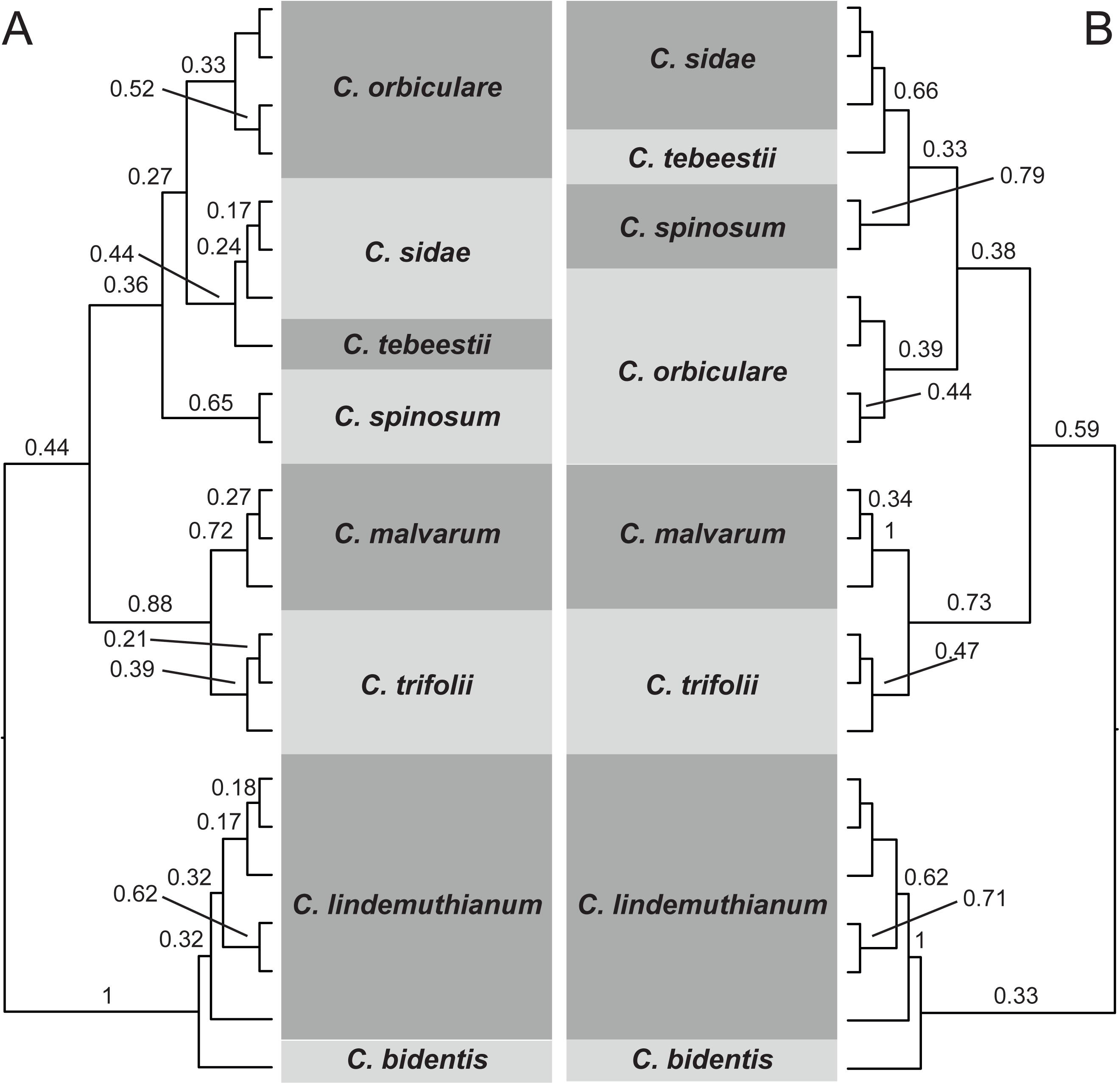
Primary concordance trees resulting from the Bayesian concordance analyses including isolates from the *C. orbiculare* complex. A. All markers (ACT, CHS-1, GAPDH, GS, HIS3, ITS and TUB2). B. Best markers (HIS3, GAPDH and GS). Concordance factors are shown above the branches that were resolved by at least one marker (≥0.14 for all markers and ≥0.33 for the best markers).

#### 3.3.11 Colletotrichum orchidearum s. I

TUB2 and HIS3 were the most powerful genes capable of discriminating species within the *C. orchidearum* complex (PIV=17 and 16 respectively), followed by CHS-1 (PIV=12) (Fig. 2). TUB2 and HIS3 are good markers to discriminate all species in the complex and support the relationships among them (PImax=0.8 and 0.67 respectively), whereas CHS-1 can help to distinguish and support recently diverged species (PImax=0.47). All species were recovered with high levels of monophyly with TUB2 (GSI=0.99—1), and also by both multilocus analyses (GSI=0.97—1) (Fig. 3). *Colletotrichum musicola* was the only species that could be recovered by all markers evaluated (CF=1) (Fig. 13). The CFs increased when the less informative markers were removed from the analysis and most species were recovered by all markers (CF=1), with the exception of *C. cliviicola and C. plurivorum* (CF≥0.66). Damm et al. (2019) also reported that some clades were not supported by some of the individual gene analyses. All deep and shallow nodes retained high support when only the best markers were combined in our analyses (Supplementary File S12B).

**Fig. 13.**
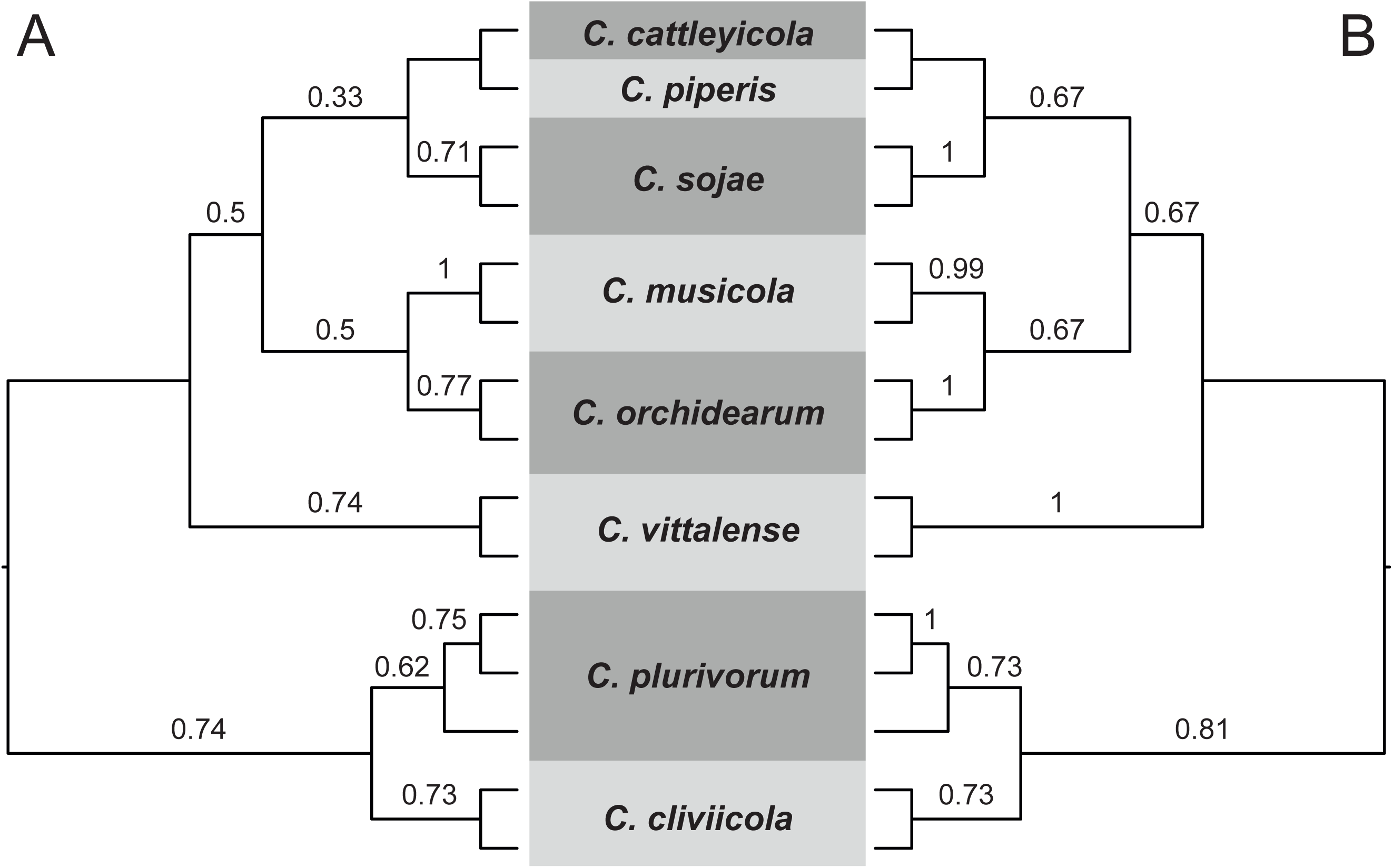
Primary concordance trees resulting from the Bayesian concordance analyses including isolates from the *C. orchidearum* complex. A. All markers (ACT, CHS-1, GAPDH, HIS3, ITS and TUB2). B. Best markers (CHS-1, HIS3 and TUB2). Concordance factors are shown above the branches that were resolved by at least one marker (≥0.16 for all markers and ≥0.33 for the best markers).

#### 3.3.12 Colletotrichum spaethianum s. I

GAPDH, TUB2 and ACT were the best markers to separate species within the *C. spaethianum* complex (Fig. 2). These markers peak at approximately 0.4 (PImax 0.47—0.77), which is in the range of where most species in the complex diverge indicating they can separate the majority of species in this complex. However, GADPH is the only marker that recovers all species as monophyletic (GSI 0.98—1) (Fig. 3). All species reached complete monophyly (GSI=1) in both multilocus trees. *Colletotrichum liriopes* and *C. spaethianum* could be recovered by most of the genes only when the best markers are considered in the BCA (CF=0.77 and 0.70, respectively) (Fig. 14B). All species remain strongly supported when only the best markers were concatenated (Supplementary Figure S13B).

**Fig. 14.**
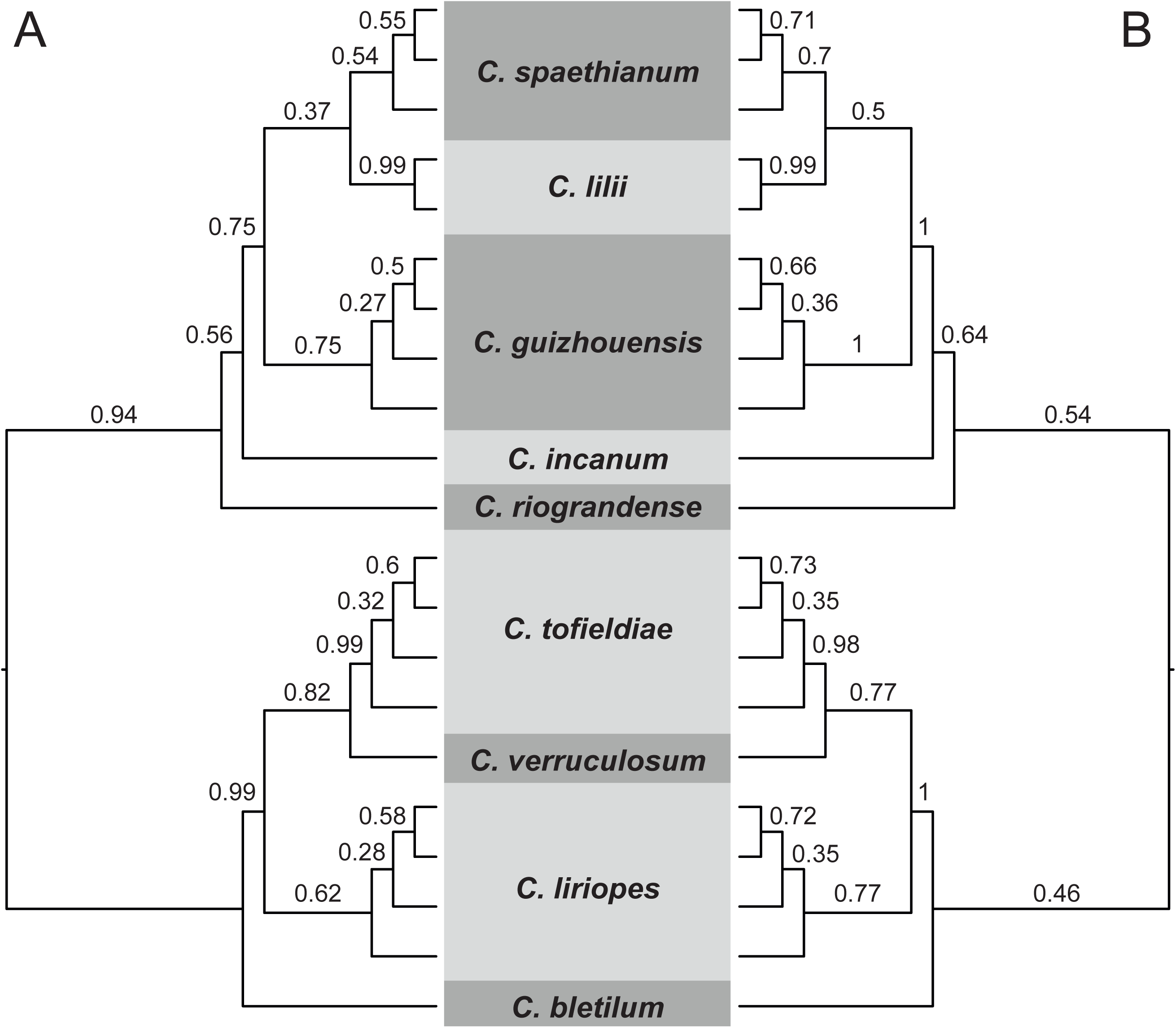
Primary concordance trees resulting from the Bayesian concordance analyses including isolates from the *C. spaethianum* complex. A. All markers (ACT, GAPDH, ITS and TUB2). B. Best markers (ACT, GAPDH, and TUB2). Concordance factors are shown above the branches that were resolved by at least one marker (≥0.25 for all markers and ≥0.33 for the best markers).

HIS3 appears to be a good marker in several other *Colletotrichum* complexes according to the present study, and may be among the three best markers for *C. spaethianum* complex. However, sequences of HIS3 and other markers such as CHS-1 and CAL are available only for one isolate of several species and we cannot include them in our analyses. Sequences of these markers need to be generated for other isolates to determine if one of these markers could be used to substitute for ACT, which does not perform well for separating *Colletotrichum* species.

#### 3.3.13 Colletotrichum truncatum s. I

GAPDH was clearly the most informative marker (PIV=57), followed by TUB2 and ACT (PIV=33 and 23 respectively) (Fig. 2). All markers peaked above 0.4 and were able to discriminate most species. *Colletotrichum acidae* and *C. curcumae* were monophyletic (GSI=1) in all single and multilocus datasets (Fig. 3) with CFs equal to 1 in analyses of both all and best markers BCA (Fig. 15). These species were also supported by maximum bootstrap values in the multilocus trees (Supplementary File S14). *Colletotrichum corchorum-capsularis* and *C. truncatum* were not recovered as monophyletic by any dataset. Moreover, *C. truncatum* is paraphyletic or polyphyletic in the multilocus trees (Fig. 15, Supplementary File S14), which leads us to the conclusion that *C. corchorum-capsularis*, as circumscribed by Niu et al. (2016), cannot be recognized as a species distinct from *C. truncatum*. Our results strongly suggest that *C. corchorum-capsularis* and *C. truncatum* may be the same species and isolates of both species were placed together in a clade with high CF (≥0.96) in the BCAs and maximum support in the multilocus analyses (Fig. 15, Supplementary File S14). Additional work is needed using the best markers for the *C. truncatum* complex and objective species recognition methods to determine the taxonomic status species boundaries of *C. corchorum-capsularis* and *C. truncatum*.

**Fig. 15.**
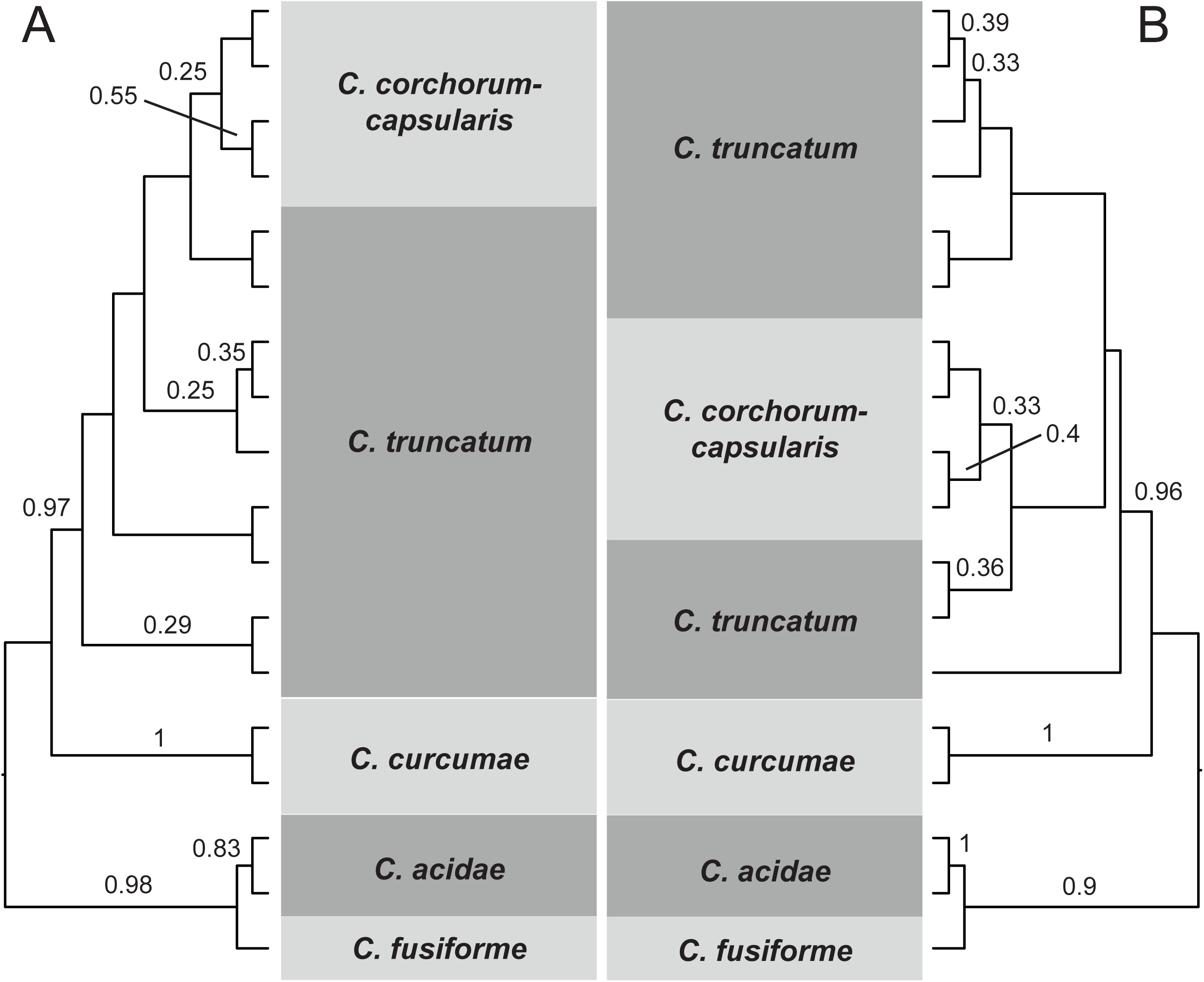
Primary concordance trees resulting from the Bayesian concordance analyses including isolates from the *C. truncatum* complex. A. All markers (ACT, GAPDH, ITS and TUB2). B. Best markers (ACT, GAPDH, and TUB2). Concordance factors are shown above the branches that were resolved by at least one marker (≥0.25 for all markers and ≥0.33 for the best markers).

## 4. Conclusions

We used phylogenetic informativeness profiling, maximum likelihood and coalescent-based phylogenetic analyses, measures of barcode utility, and genealogical sorting indices to assess the performance of the several molecular markers used in *Colletotrichum* systematics and taxonomy across all known species complexes. While HIS3, GAPDH, and TUB2 were among the best markers for most of the complexes, the optimal set of markers is not always the same across all complexes. ACT, CHS-1 and nrITS were the worst markers and, as previously proposed for the *C. gloeosporioides* complex (Vieira et al., 2017), they can be discarded from the phylogenetic analysis of almost all species complexes. ACT was retained in the set of best markers for the *C. graminicola*, *C. spaethianum* and *C. truncatum* complexes to achieve a minimum of three markers as proposed in the methodology of the present study. However, few markers were included for these complexes due to missing data, therefore additional markers need to be sequenced and their performance evaluated. The analyses of *C. caudatum, C. dematium, C. graminicola* and *C. spaethianum* complexes were the most impacted by the excessive amount of missing data for the majority of the markers, which highlights the importance of selecting a standard set of markers to delimit species. Similarly, several isolates and/or species were not included in the marker analyses for *C. gloeosporioides s.l.* due to selective data acquisition by different research groups. It is not clear how the inclusion of sequences from these isolates might impact our results. Sequences need to be generated for these markers and/or species to provide more decisive results. We have also identified species complexes, which will need to be revisited in the future, in which it appears species have been misidentified (*C. acutatum*, *C. dematium* and *C. truncatum)*.

Selecting the optimal markers to sequence for biodiversity studies on *Colletotrichum* will impact *Colletotrichum* studies in a few ways. First, species recognition will likely be more accurate and robust by avoiding the confounding effect of including markers with low phylogenetic signal in the analyses. Secondly, phylogenetic studies of *Colletotrichum* will become more economical, since sequencing markers with low informativeness represents a low return on investment. Finally, if research groups take guidance from this study, we are more likely to see a consensus developed on the data acquired for phylogenetic studies on *Colleotrichum* and we will be closer to a global assessment with combinable data.

Researchers around the world continue to have an interest in documenting the diversity of *Colletotrichum* species associated with economically important plant species. However, this work is labor intensive and expensive because several hundred isolates are typically screened and a paucity of distinctive morphological characters necessitates DNA sequencing. The expense has been unnecessarily compounded by the lack of an objective and comprehensive assessment of the utility of existing markers for phylogenetic inference and species identification/delimitation and the lack of a consensus on the markers to be used. We hope the results presented here will help to address this problem. While the optimal markers differ by species complex, our results provide some guidance on the most efficient path to document and describe diversity within *Colletotrichum*. Our results also show that for the accurate identification and delimitation of *Colletotrichum* species, a small set of markers with strong phylogenetic signal is more suitable than a large set including markers with both weak and strong phylogenetic signal. GAPDH is among the optimal set of markers for 10 of the 13 species complexes in *Colletotrichum*, followed by TUB2 (10 of 13), and HIS3 (7 of 13). Therefore, GAPDH is a good marker to sequence for initial diversity screening and assigning isolates to a species complex because data for this marker is available for the majority of species within the genus. However, selection of additional markers for phylogenetic inference and species delimitation will depend on the species complex.

Finally, while we recommend the optimal markers for species recognition within *Colletotrichum* in order to improve diversity studies in the genus, our understanding of evolutionary relationships among species remains poorly resolved. Improving our understanding of relationships among taxa within *Colletotrichum* will require more robust genomic sampling. Genome sequencing is underway for many species of *Colletotrichum*, however a comprehensive phylogenomic study of the genus is needed.

## Acknowledgments

This study was funded by “Conselho Nacional de Desenvolvimento Científico e Tecnológico – CNPq” (Universal number 408724/2018-8). Willie A. S. Vieira acknowledges the “Fundação de Amparo à Ciência e Tecnologia do Estado de Pernambuco – FACEPE” for the Postdoc fellowship (Process number BFP-0040-5.01/16). Marcos P. S. Câmara acknowledge CNPq for the research productivity fellows. Josiene S. Veloso acknowledge the “Coordenação de Aperfeiçoamento Pessoal de Ensino Superior – CAPES” and the “Programa Nacional de Pós-Doutorado/CAPES – PNPD/CAPES” for the Posdoc fellowship. This work represents partial fulfillment of the degree of Master of Phytopathology for Priscila A. Bezerra. V. P. Doyle would like to acknowledge funding support from the Louisiana Board of Regents Research Competitiveness Subprogram (LEQSF(2016-19)-RD-A-01).

## Glossary

**Appressorium**: specialized cell produced by some phytopathogenic fungi which is used to infect plant hosts.

**Conidium**: asexual spore of Ascomycota and Basidiomycota.

**Endophytic fungi**: fungi that grow inside the plant tissues without causing disease symptoms.

**Phytopathogenic fungi**: fungi that cause plant diseases.

***Sensu lato* (*s. l.*)**: taxonomic terminology used to reference species complexes (*C. acutatum* species complex = *C. acutatum s. l.*).

***Sensu stricto* (*s. s.*)**: when is necessary to refer the species with the same name of the complex (*C. acutatum s. s.* is a species within *C. acutatum s. l.*).

**Species complex**: major clades strongly supported within *Colletotrichum* genus tree. These clades include phylogenetic species closely related which most are indistinguishable based on phenotypical characters (e.g. conidial and appressorial shape and size, growth rate, color of colonies). Species complexes get the same name of the species within them that is more known or that was firstly described. In some cases, members within a given species complex share peculiar conidial characteristics: *C. acutatum* – conidia with acute ends; *C. boninense* – presence of a prominent scar (hilum) at the base of the conidium; *C. caudatum* – conidia with a filiform appendage at the apex; *C. gigasporum* – longest and widest conidia within the genus.

## References

Ané, C., Larget, B., Baum, D.A., Smith, S.D., Rokas, A., 2007. Bayesian estimation of concordance among gene trees. Molecular Biology and Evolution 24, 412–426.

Babicki, S., Arndt, D., Marcu, A., Liang, Y., Grant, J.R., Maciejewski, A., Wishart, D.S., 2016. Heatmapper: web-enabled heat mapping for all. Nucleic Acids Research 44, W147–W153.

Baum, D.A., 2007. Concordance trees, concordance factors, and the exploration of reticulate genealogy. Taxon 56, 417–426.

Bragança, C.A.D., Damm, U., Baroncelli, R., Massola Júnior, N.S., Crous, P.W., 2016. Species of the *Colletotrichum acutatum* complex associated with anthracnose diseases of fruit in Brazil. Fungal Biology 120, 547–561.

Cai, L., Hyde, K.D., Taylor, P.W.J., Weir, B.S., Waller, J., Abang, M.M., Zhang, J.Z., Yang, Y.L., Phoulivong, S., Liu, Z.Y., Prihastuti, H., Shivas, R.G., McKenzie, E.H.C., Johnston, P.R., 2009. A polyphasic approach for studying *Colletotrichum*. Fungal Diversity 39, 183–124.

Cannon, P.F., Damm, U., Johnston, P., Weir, B.S., 2012. *Colletotrichum* – Current status and future directions. Studies in Mycology 73, 181–213.

Cao, X.; Xu, X.; Che, H.; West, J.S.; Luo, D., 2019. Three Colletotrichum Species, Including a New Species, are Associated to Leaf Anthracnose of Rubber Tree in Hainan, China. Plant Disease 103, 117–124.

Corda, A.C.I., 1831. Die Pilze Deutschlands. In: Deutschlands Flora in Abbildungen nach der Natur mit Beschreibungen (Sturm, J, ed.). Sturm, Nürnberg vol. 3, Abt. 12: 33–64, tab. 21–32.

Costa, J.F.O., Kamei, S.H., Silva, J.R.A., Miranada, A.R.G.S., Netto, M.B., Silva, S.J.C., Correia, K.C., Lima, G.S.A., Assunção, I.P., 2018. Species diversity of *Colletotrichum* infecting *Annona* spp. in Brazil. European Journal of Plant Pathology. https://doi.org/10.1007/s10658-018-01630-w.

Crouch, J.A., Clarke, B.B., Hillman, B.I, 2009a. What is the value of ITS sequence data in *Colletotrichum* systematics and species diagnosis? A case study using the falcate-spored graminicolous *Colletotrichum* group. Mycologia 101, 648–656.

Crouch, J.A., Beirn, L.A., Cortese, L.M., Bonos, S.A., Clarke, B.B., 2009b. Anthracnose disease of switchgrass caused by the novel fungal species *Colletotrichum navitas*. Mycological Research 113, 1411–1421

Crouch, J.A., Tredway, L.P., Clarke, B.B. and Hillman, B.I., 2009c. Phylogenetic and population genetic divergence correspond with habitat for the pathogen *Colletotrichum cereale* and allied taxa across diverse grass communities. Molecular Ecology 18, 123–135.

Crouch, J.A., Tomaso-Peterson, M., 2012. Anthracnose disease of centipedegrass turf caused by *Colletotrichum eremochloae*, a new fungal species closely related to *Colletotrichum sublineola*. Mycologia 104, 1085–1096.

Cummings, M.P., Neel, M.C., Shaw, K.L., 2008. A genealogical approach to quantifying lineage divergence. Evolution 62, 2411–2422.

Damm, U., Woudenberg, J.H.C., Cannon, P.F., Crous, P.W., 2009. *Colletotrichum* species with curved conidia from herbaceous hosts. Fungal Diversity 39, 45–87.

Damm, U., Cannon, P.F., Liu, F., Barreto, R.W., Guatimosim, E, Crous, P.W., 2013. The *Colletotrichum orbiculare* species complex: Important pathogens of field crops and weeds. Fungal Diversity 61, 29–59.

Damm, U., Cannon, P.F., Woudenberg, J.H.C., Crous, P., 2012a. The *Colletotrichum acutatum* species complex. Studies in Mycology 73, 37–113.

Damm, U., Cannon, P.F., Woudenberg, J.H.C., Johnston, P.R., Weir, B.S., Tan, Y.P., Shivas, R.G., Crous, P.W., 2012b. The *Colletotrichum boninense* species complex. Studies in Mycology 73:1–36.

Damm, U., Sato, T., Alizadeh, A., Groenewald, J.Z., Crous, P.W., 2019. The *Colletotrichum dracaenophilum*, C. magnum and C. orchidearum species complexes. Studies in Mycology 92, 1–46.

Dean, R., Van Kan, J.A.L., Pretorius, Z.A., Hammond-Kosack, K.E., Di Pietro, A., Spanu, P.D., Rudd, J.J., Dickman, M., Kahmann, R., Ellis, J., Foster, G.D., 2012. The Top 10 fungal pathogens in molecular plant pathology. Molecular Plant Pathology 13, 414–430.

Dettman, J.R., Jacobson, D.J., Taylor, J.W., 2003. A multilocus genealogical approach to phylogenetic species recognition in the model eukaryote *Neurospora*. Evolution 57, 2703–2720.

Diao, Y.-Z., Zhang, C., Liu, F., Wang, W.-Z., Liu, L., Cai, L., Liu, X.-L., 2017. *Colletotrichum* species causing anthracnose disease of chili in China. Persoonia 38, 20–37.

Doyle, V.P., Oudemans, P.V., Rehner, S.A., Litt, A., 2013. Habitat and host indicate lineage identity in *Colletotrichum gloeosporioides s. l.* from wild and agricultural landscapes in North America. PLoS ONE 8, e62394.

Du, M., Schardl, C., Nuckles, E., Vaillancourt, L., 2005. Using mating-type gene sequences for improved phylogenetic resolution of *Collectotrichum* species complexes. Mycologia 97, 641–658.

Fong, J.J., Fujita, M.K., 2011. Evaluating phylogenetic informativeness and data-type usage for new protein-coding genes across Vertebrata. Molecular Phylogenetics and Evolution 61, 300–307.

Fu, M., Crous, P.W., Bai, Q., Zhang, P.F., Xiang, J., Guo, Y.S., Zhao, F.F., Yang, M.M., Hong, N., Xu, W.X. and Wang, G.P., 2019. *Colletotrichum* species associated with anthracnose of *Pyrus* spp. in China. Persoonia 42, 1–35.

Hebert, P.D., Cywinska, A., Ball, S.L., 2003. Biological identifications through DNA barcodes. Proceedings of the Royal Society of London Series B 270, 313–321.

Hyde KD, Cai L, McKenzie EHC, Yang YL, Zhang JZ, Prihastuti, H., 2009. *Colletotrichum*: a catalogue of confusion. Fungal Diversity 39, 1–17.

Hyde, K.D., Udayanga, D., Manamgoda, D.S., Tedersoo, L., Larsson, E., Abarenkov K., Bertrand, Y.J.K., Oxelman, B., Hartmann, M., Kauserud, H., Ryberg, M., Kristiansson, E., Nilsson, R.H., 2013. Incorporating molecular data in fungal systematics: A guide for aspiring researchers. Current Research in Environmental and Applied Mycology 3, 1–32.

Jayawardena, R.S., Yan, J., Hyde, K.D., Zhang, G.,2016. Morphological and molecular characterization of *Colletotrichum* species of strawberry in China. Mycosphere 7, 1147–1163.

Katoh, K., Misawa, K., Kuma, K., Miyata, T., 2002. MAFFT: a novel method for rapid multiple sequence alignment based on fast Fourier transform. Nucleic Acids Research 30, 3059–3066.

Katoh, K., Standley, D.M, 2013. MAFFT multiple sequence alignment software version 7: Improvements in performance and usability. Molecular Biology and Evolution 30, 772–780.

Kumar, S., Stecher, G., Tamura, K., 2016. MEGA7: Molecular Evolutionary Genetics Analysis version 7.0 for bigger datasets. Molecular Biology and Evolution 33, 1870–1874.

Larget, B., Kotha, S.K., Dewey, C.N., Ané, C., 2010. BUCKy: Gene tree/species tree reconciliation with the Bayesian concordance analysis. Bioinformatics 26, 2910–2911.

Librado, P., Rozas, J.,2009. DnaSP v5: a software for comprehensive analysis of DNA polymorphism data. Bioinformatics 25, 1451–1452.

Lima, N.B., Batista, M.V.A., Morais Jr, M.A., Barbosa, M.A.G., Michereff, S.J., Hyde, K.D., Câmara, M.P.S., 2013. Five *Colletotrichum* species are responsible for mango anthracnose in northeastern Brazil. Fungal Diversity 61:75–88.

Liu, F., Cai, L., Crous, P.W., Damm, U., 2014. The *Colletotrichum gigasporum* species complex. Persoonia 33, 83–97.

Liu, F., Wang, M., Damm, U., Crous, P.W., Cai, L., 2016. Species boundaries in plant pathogenic fungi: A *Colletotrichum* case study. BMC Evolutionary Biology 16, 81.

Liu, F., Weir, B.S., Damm, U., Crous, P.W., Wang, Y., Liu, B., Wang, M., Zhang, M., Cai, L., 2015. Unravelling *Colletotrichum* species associated with *Camellia*: Employing ApMat and GS loci to resolve species in the *C. gloeosporioides* complex. Persoonia 35, 63–86.

Lopez-Giraldez, F., Townsend, J.P., 2011. PhyDesign: An online application for profiling phylogenetic informativeness. BMC Evolutionary Biology 11, 152.

Magain, N., Miadlikowska, J., Mueller, O., Gajdeczka, M., Truong, C., Salamov, A.A., Dubchak, I., Grigoriev, I.V., Goffinet, B., Sérusiaux, E., Lutzoni, F., 2017. Conserved genomic collinearity as a source of broadly applicable, fast evolving, markers to resolve species complexes: A case study using the lichen-forming genus Peltigera section Polydactylon. Molecular Phylogenetics and Evolution 117, 10–29.

Marin-Felix, Y., Groenewald, J.Z., Cai, L., Chen, Q., Marincowitz, S., Barnes, I., Bensch, K., Braun, U., Camporesi, E., Damm, U., de Beer, Z.W., Dissanayake, A., Edwards, J., Giraldo, A., Hernandez-Restrepo, M., Hyde, K.D., Jayawardena, R.S., Lombard, L., Luangsa-Ard, J., McTaggart, A.R., Rossman, A.Y., Sandoval-Denis, M., Shen, M., Shivas, R.G., Tan, Y.P., van der Linde, E.J., Wingfield, M.J., Wood, A.R., Zhang, J.Q., Zhang, Y., Crous, P.W., 2017. Genera of phytopathogenic fungi: GOPHY 1. Studies in Mycology 86, 99–216.

Mills, P.R., Hodson, A., Brown, A.E., 1992. Molecular differentiation of Colletotrichum gloeosporioides isolates infecting tropical fruits, in: Bailey, J.A., Jeger, M.J. (Eds.), Colletotrichum: Biology, Pathology and Control, CABI, Wallingford, 269–288.

Moriwaki, J., Tsukiboshi, T., 2009. *Colletotrichum echinochloae*, a new species on Japanese barnyard millet (*Echinochloa utilis*). Mycoscience 50, 273–280.

Niu, X., Gao, H., Qi, J., Chen, M., Tao, A., Xu, J., Dai, Z., Su, J., 2016. Colletotrichum species associated with jute (Corchorus capsularis L.) anthracnose in southeastern China. Scientific Reports 6, 25179.

O’Connell, R.J., Thon, M.R., Hacquard, S., Amyotte, S.G., Kleemann, J., Torres, M.F., Damm, U., Buiate, E.A., Epstein, L., Alkan, N., Altmuller, J., Alvarado-Balderrama, L., Bauser, C.A., Becker, C., Birren, B.W., Chen, Z., Choi, J., Crouch, J.A., Duvick, J.P., Farman, M.A., Gan, P., Heiman, D., Henrissat, B., Howard, R.J., Kabbage, M., Koch, C., Kracher, B., Kubo, Y., Law, A.D., Lebrun, M.H., Lee, Y.H., Miyara, I., Moore, N., Neumann, U., Nordstrom, K., Panaccione, D.G., Panstruga, R., Place, M., Proctor, R.H., Prusky, D., Rech, G., Reinhardt, R., Rollins, J.A., Rounsley, S., Schardl, C.L., Schwartz, D.C., Shenoy, N., Shirasu, K., Sikhakolli, U.R., Stuber, K., Sukno, S.A., Sweigard, J.A., Takano, Y., Takahara, H., Trail, F., Van Der Does, H.C., Voll, L.M., Will, I., Young, S., Zeng, Q., Zhang, J., Zhou, S., Dickman, M.B., Schulze-Lefert, P., Van Themaat, E.V., Ma, L.J., Vaillancourt, L.J., 2012. Lifestyle transitions in plant pathogenic *Colletotrichum* fungi deciphered by genome and transcriptome analyses. Nature Genetics 44, 1060–1065.

Oliveira, L.F.M., Feijó, F.M., Mendes, A.L.S.F., Neto, J.D.V., Netto, M.S.B., Assunção, I.P., Lima, G.S.A., 2018. Identification of *Colletotrichum* species associated with brown spot of cactus prickly pear in Brazil. Tropicala Plant Pathology 43, 247–253.

Paradis, E., Claude, J., Strimmer, K., 2004. APE: Analyses of phylogenetics and evolution in R language. Bioinformatics 20, 289–290.

Pond, S.L., Frost, S.D., Muse, S.V., 2005. HyPhy: hypothesis testing using phylogenies. Bioinformatics 21, 676–679.

R Core Team. 2017. R: A language and environment for statistical computing. R Foundation for Statistical Computing, Vienna, Austria.

Rojas, E.I., Rehner, S.A., Samuels, G.J., Van Bael, S.A., Herre, E.A., Cannon, P., Chen, R., Pang, J., Wang, R., Zhang, Y., Peng, Y.Q., Sha, T., 2010. *Colletotrichum gloeosporioides s.l.* associated with *Theobroma cacao* and other plants in Panamá: Multilocus phylogenies distinguish host-associated pathogens from asymptomatic endophytes. Mycologia 102,1318–1338.

Ronquist, F., Teslenko, M., van der Mark, P., Ayres, D.L., Darling, A., Höhna, S., Larget, B., Liu, L., Suchard, M.A., Huelsenbeck, J.P., 2012. MrBayes v. 3.2: Efficient Bayesian phylogenetic inference and model choice across a large model space. Systematic Biology 61, 539–542.

Sakalidis, M.L., Hardy, G.E.S.J., Burgess, T.I., 2011. Use of the Genealogical Sorting Index (GSI) to delineate species boundaries in the *Neofusicoccum parvum-Neofusicoccum ribis* species complex. Molecular Phylogenetics and Evolution, 60, 333–344.

Samarakoon, M.C., Peršoh, D., Hyde, K.D., Bulgakov, T.S., Manawasinghe, I.S., Jayawardena, R.S., Promputtha, I., 2018. *Colletotrichum acidae* sp. nov. from northern Thailand and a new record of *C. dematium* on Iris sp. Mycosphere 9: 583–597.

Schmitt,.I, Crespo, A., Divakar, P.K., Fankhauser, J.D., Herman-Sackett, E., Kalb, K., Nelsen, M.P., Nelson, N.A., Rivas-Plata, E., Shimp, A.D.,Widhelm, T., Lumbsch, H.T., 2009. New primers for promising single-copy genes in fungal phylogenetics and systematics. Persoonia 23, 35–40.

Sela, I., Ashkenazy, H., Katoh, K., Pupko, T., 2015. GUIDANCE2: Accurate detection of unreliable alignment regions accounting for the uncertainty of multiple parameters. Nucleic Acids Research 43, W7–W14.

Sharma, G., Kumar, N., Weir, B.S., Hyde, K.D., Shenoy, B.D., 2013. The ApMat marker can resolve *Colletotrichum* species: A case study with *Mangifera indica*. Fungal Diversity 61,117–38.

Sharma, G., Pinnaka, A.K., Belle, D.S., 2015. Resolving the *Colletotrichum siamense* species complex using ApMat marker. Fungal Diversity 71, 247–64.

Sharma, G., Maymon, M., Freeman, S., 2017. Epidemiology, pathology and identification of Colletotrichum including a novel species associated with avocado (Persea americana) anthracnose in Israel. Scientific Reports 7, 15839.

Silva, D.N., Talhinas, P., Várzea, V., Cai, L., Paulo, O.S., Batista, D., 2012. Application of the Apn2/MAT locus to improve the systematics of the *Colletotrichum gloeosporioides* complex: An example from coffee (*Coffea* spp.) hosts. Mycologia 104, 396–409.

Silva, J.R.A, Chaves, T.P., Silva, a A.R.G., Barbosa, L.F., Costa, F.F.O., Ramos-Sobrinho, R., Teixeira, R.R.O., Silva, S.J.C., Lima, G.S.A., Assunção, I.P., 2017. Molecular and morpho-cultural characterization of *Colletotrichum* spp. associated with anthracnose on *Capsicum* spp. in northeastern Brazil. Tropical Plant Pathology 42, 315–319.

Sousa, E.S., Silva, J.R.A., Assunção, I.P., Melo, M.P., Feijó, F.M., Matos, K.S., Lima, G.S.A., Beserra Jr, J.E.A., 2018. *Colletotrichum* species causing anthracnose on lima bean in Brazil. Tropical Plant Pathology 43, 78–84.

Sreenivasaprasad, S., Brown, A.E., Mills, P.R., 1992. DNA sequence variation and interrelationship among *Colletotrichum* species causing strawberry anthracnose. Physiological and Molecular Plant Pathology 41, 265–281.

Stawatakis, A., 2014. RAxML version 8: A tool for Phylogenetic Analysis and Post-Analysis of Large Phylogenies. Bioinformatics 30, 1312–1313.

Sutton, B.C., 1980. The Coelomycetes. Fungi Imperfecti with Pycnidia, Acervuli and Stromata. CABI, Kew

Tao, G., Liu, Z.Y., Liu, F., Gao, Y.H., Cai, L., 2013. Endophytic Colletotrichum species from Bletilla ochracea (Orchidaceae), with descriptions of seven new speices. Fungal Diversity 61, 139–164.

Taylor, J.W., Jacobson, D.J., Kroken, S., Kasuga, T., Geiser, D.M., Hibbett, D.S., Fisher, M.C., 2000. Phylogenetic species recognition and species concepts in fungi. Fungal Genetics and Biology 31, 21–32.

Vaidya, G., Lohman, D. J., Meier, R., 2011. SequenceMatrix: concatenation software for the fast assembly of multi-gene datasets with character set and codon information. Cladistics 27, 171–180.

Veloso, J.S., Câmara, M.P.S., Lima, W.G., Michereff, S.J., Doyle, V.P.. (2018). Why species delimitation matters for fungal ecology: *Colletotrichum* diversity on wild and cultivated cashew in Brazil. Fungal Biology 122, 677–691.

Vieira, W.A., Michereff, S.J., de Morais Jr, M.A., Hyde, K.D., Câmara, M.P.S., 2014. Endophytic species of *Colletotrichum* associated with mango in northeastern Brazil. Fungal Diversity 67, 181–202.

Vieira, W.A.S., Lima, W.G., Nascimento, E.S., Michereff, S.J., Câmara, M.P.S., Doyle, V.P., 2017. The impact of phenotypic and molecular data on the inference of *Colletotrichum* diversity associated with *Musa*. Mycologia 109, 912–934.

von Arx, J.A., 1957. Die Arten der Gattung Colletotrichum Cda. Phytopathologische Zeitschrift 29, 413–468.

Wang, Q.T., Liu, X.T., Ma, H.Y., Shen, X.Y., Hou, C.L., 2019. *Colletotrichum yulongense sp. nov.* and *C. rhombiforme* isolated as endophytes from *Vaccinium dunalianum var. urophyllum* in China. Phytotaxa 394, 285–298.

Weir, B.S., Johnston, P.R., Damm, U., 2012. The *Colletotrichum gloeosporioides* species complex. Studies in Mycology 73,115–180.

